# Jlp2 is a non-canonical release factor that protects cells during ribosome stalling

**DOI:** 10.1101/2025.09.04.673968

**Authors:** Kaushik Viswanathan Iyer, Alina-Andrea Kraft, Chloé Astrid Walter, Max Müller, Lena Sophie Tittel, Grit Florentine Stampa, Marie-Luise Winz

## Abstract

Ribosome stalling generates aberrant nascent peptides that remain tethered to the large ribosomal subunit following ribosome splitting. Such peptidyl-tRNA:60S complexes are processed by the Ribosome-associated Quality Control (RQC) pathway, where Ltn1 and Rqc1 mediate K48-linked ubiquitination and Rqc2 adds CAT tails to the nascent peptides. Subsequently, Vms1 releases the peptides from tRNA, enabling their proteasomal degradation. However, alternative mechanisms that process these stalled intermediates remain poorly defined. In this study, we identify Jlp2 as a novel factor involved in translation quality control. We find that Jlp2 is a release factor that catalyzes peptide release from tRNA and suppresses excessive CAT tailing when Ltn1-dependent ubiquitination is compromised. We define its ribosome binding properties, substrate scope, and critical residues required for peptide release. Our data support a model in which Jlp2-mediated peptide release constitutes an alternative quality control mechanism to safeguard cells during ribosome stalling and conditions of RQC failure or insufficiency.

## Introduction

Despite the high ribosomal translation fidelity, ribosome stalling frequently occurs due to mRNA damage or structure, aminoacyl-tRNA scarcity or ‘problematic’ codon or amino acid sequences^1–8^. Ribosomal stalling on actively translated mRNA often results in ribosome collisions^9–11^, presenting several points of concern for resolution by the cell, including ribosome recycling to maintain translation capacity, and clearance of mRNA and nascent peptides associated with collisions to prevent cellular accumulation of potentially toxic molecules.

A highly conserved network of co-Translational Quality Control (TQC) pathways resolves ribosome collisions in eukaryotes^8,12^. In yeast, Hel2 (ZNF598 in mammals) binds collided disomes and initiates the surveillance response by K63-linked polyubiquitination of ribosomal protein uS10^9,13–15^, triggering ribosome splitting by the RQT complex^14,16,17^. Alternatively, Dom34 (PELOTA in mammals) with cofactors Hbs1 and Rli1 (ABCE1 in mammals) can split ribosomes^18–22^ independent of Hel2. Critically, these unconventional termination mechanisms leave 60S ribosomal subunits occupied by nascent peptidyl-tRNA due to lack of peptidyl-tRNA hydrolysis^16,23–26^.

Peptidyl-tRNA:60S complexes become substrates for the Ribosome-associated Quality Control (RQC) complex composed of Ltn1, Rqc1 and Rqc2 (LISTERIN, TCF25 and NEMF in mammals, respectively)^24,27–30^. The E3 ubiquitin ligase Ltn1, with Rqc1, polyubiquitinates nascent peptide lysines in a K48-linked manner, triggering proteasomal degradation^24,28,31–34^. Rqc2 binds to the 60S intersubunit interface, facilitates Ltn1 recruitment^29,32,35^ and catalyzes an mRNA-independent addition of C-terminal Alanine and Threonine (CAT) tails to nascent peptides^30,36^. CAT tails evict lysines hidden in the peptide exit tunnel and promote CAT tail-dependent degradation^37–39^. However, excessive CAT tailing without Ltn1-mediated ubiquitination results in CAT tail-dependent, proteasome-resistant aggregates^40^, highlighting a pathological consequence of RQC imbalance. Ubiquitinated nascent peptides are extracted by the Cdc48 (VCP/p97 in mammals)-Ufd1-Npl4 complex and delivered to the proteasome^32–34^. Extraction requires peptide release from tRNA, catalyzed by Vms1 (ANKZF1 in mammals) and its co-factor Arb1, via endonucleolytic tRNA cleavage near the 3’-end^28,41–43^. Additionally, Vms1 counteracts CAT tailing by competing with Rqc2 for 60S binding^44^.

While canonical RQC provides an elegant multi-step resolution of peptidyl-tRNA:60S complexes, this process requires the coordinated activity of numerous proteins. Consequently, failure at any step may result in the disruption of the entire cascade. This raises a fundamental question — are there alternative mechanisms that can process stalled intermediates when the canonical pathway fails?

In this study, we identify the uncharacterized *Saccharomyces cerevisiae* protein Jlp2 as a novel quality control factor that resolves substrates arising from ribosome stalling. We show that *JLP2* protects cells during ribosome stalling stress by catalyzing peptide release from tRNA, thereby suppressing excessive CAT tailing and CAT tail-dependent aggregation in the absence of Ltn1-dependent ubiquitination. Importantly, this function is independent of the RQC complex in addition to several known TQC release factors. Together, our findings define an alternative TQC pathway and the critical role of Jlp2 in safeguarding proteostasis when canonical quality control is compromised.

## Results

### Jlp2 chemogenetically interacts with members of the RQC complex

To identify novel factors involved in TQC, we screened publicly available high-throughput genetic, physical and chemogenetic interaction databases for candidates showing interactions and/or correlations to known TQC components^45–50^. This identified Jlp2, which showed strong chemogenetic correlations to many TQC factors, with its deletion causing among the strongest sensitivities to the translation elongation inhibitors CHX and ANI^46,50^. This suggested that Jlp2 may play an important role during translation elongation stress, particularly associated with ribosome collisions. Notably, Jlp2 is one of only two yeast proteins containing an NFACT RNA-binding (NFACT-R) domain^51^ (Fig. 1a), the other being Rqc2 - a core RQC component, further supporting a potential function in TQC.

**Figure 1.**
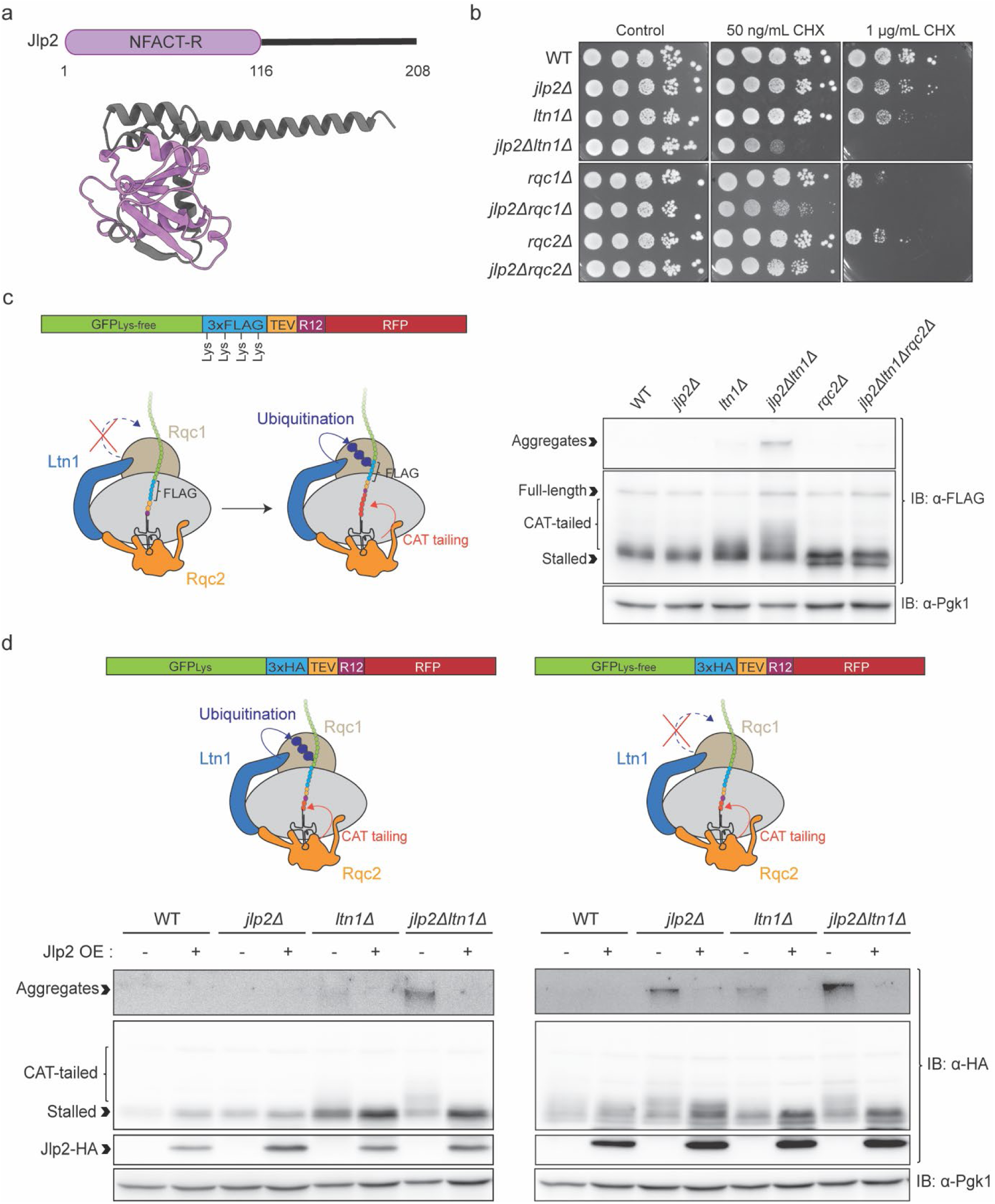
Jlp2 is involved in the quality control of substrates arising from ribosome collisions. a. Schematic overview of the primary structure of Jlp2 (top) and the corresponding color-matched AlphaFold-predicted secondary structure (bottom) [AF-P40206-F1-v6 (average pLDDT= 94.69)]^52^. b. *JLP2* genetically interacts with members of the RQC complex. Growth of WT, *jlp2Δ, ltn1Δ, jlp2Δltn1Δ, rqc1Δ, jlp2Δrqc1Δ, rqc2Δ, jlp2Δrqc2Δ* cells spotted in serial dilutions on YPD control and CHX-containing plates, imaged after 2 days (control), 4 days (50 ng/mL), and 10 days (1 µg/mL). Growth defects result in smaller colonies relative to WT. c. Simultaneous deletion of *JLP2* and *LTN1* results in an increase in CAT tailing and CAT tail-dependent aggregates. Western blot analysis of stall-inducing reporter GFP_Lys-free_-3xFLAG-TEV-R12-RFP in WT, *jlp2Δ, ltn1Δ, jlp2Δltn1Δ, rqc2Δ* and *jlp2Δltn1Δrqc2Δ*. Reporter structure is detailed on the left showing the requirement of CAT tailing to push the first lysine residues of the reporter (from FLAG) out of the peptide exit tunnel, exposing them to Ltn1-mediated polyubiquitination. d. Jlp2 suppresses CAT tailing of a substrate that cannot be ubiquitinated by Ltn1. Western blot analysis of GFP_Lys_-3xHA-TEV-R12-RFP and GFP_Lys-free_-3xHA-TEV-R12-RFP reporters in WT, *jlp2Δ, ltn1Δ* and *jlp2Δltn1Δ* containing either an empty vector or a Jlp2-HA overexpression vector. Schematic representations of the reporter constructs and their expected outcomes are shown at the top of the corresponding panels.

To characterize the functional relationship of Jlp2 with the RQC complex, we assessed the CHX and ANI sensitivity of *JLP2* and RQC factor deletion strains at concentrations that induce translatome-wide ribosome collisions^10^. Deletion of *JLP2* alone showed a modest growth defect compared to wild type (WT) cells in the presence of 50 ng/mL CHX that was visible at the 24 hours (Supplementary Fig. S1a) and diminished with time (48 hours, Supplementary Fig. S1a and Day 4, Fig. 1b). We observed similar results in the presence of 10 µg/mL ANI (Supplementary Fig. S1a). Deletion of *JLP2* resulted in a sustained growth defect in the presence of 1 µg/mL CHX (Fig. 1b), showing that Jlp2 contributes to cellular fitness during translation elongation stress. Simultaneous deletion of *JLP2* and either *LTN1* or *RQC1* resulted in strong synthetic growth defects in the presence of 50 ng/mL CHX not observed for any of the single deletants (Fig. 1b). This was also true, though to a lesser extent, for the simultaneous deletion of *JLP2* and *RQC2,* suggesting that Jlp2 and the RQC complex may be parallelly involved in a common biological process. Particularly, the strong growth defects upon loss of Jlp2 along with Ltn1 (which catalyzes nascent peptide ubiquitination)^24,31^ and Rqc1 (which specifies K48-linkage of the ubiquitin chain)^28,33^ suggested that Jlp2 may be particularly important in the absence of K48-linked nascent chain ubiquitination during RQC, since single deletants remain relatively unaffected compared to WT at this concentration (Fig. 1b). Deletion of *JLP2* and *LTN1* in the presence of 10 µg/mL ANI recapitulated the synthetic growth defect (Supplementary Fig. S1a) supporting the notion that this defect is not CHX-specific but linked to ribosome collisions. At the higher CHX concentration (1 µg/mL), double deletion of *JLP2* along with any of the RQC complex factors completely inhibited the growth of cells (Fig. 1b), further underscoring the importance of Jlp2 during translation elongation stress in the absence of RQC complex.

### Jlp2 counteracts Rqc2-mediated CAT tailing and aggregate formation of aberrantly stalled nascent peptides that escape Ltn1-dependent RQC

Given the presence of an NFACT-R domain in both Jlp2 and Rqc2, we asked if Jlp2 plays a role in the RQC of aberrant peptides arising from ribosome stalling with particular emphasis on CAT tailing. We employed a reporter that induces ribosome collisions within the mRNA on a sequence encoding 12 arginines (R12). This reporter encodes lysine-free GFP followed by a 3xFLAG tag, a TEV protease recognition sequence, an R12 sequence, and RFP (GFP_Lys-free_-3xFLAG-TEV-R12-RFP)^37^. Ribosome stalling on this reporter positions the first translated lysines (in the 3xFLAG sequence) inside the peptide exit tunnel. Hence, for Ltn1-mediated ubiquitination, CAT tailing is required to push these lysines out of the ribosome (Fig. 1c)^37^. As expected, in the WT strain, most of the stalled reporter protein was degraded following Ltn1-mediated ubiquitination (Fig. 1c). Absence of *JLP2* alone did not affect the stalled reporter peptide showing that Jlp2 was not required for CAT tailing. Absence of *LTN1* alone stabilized the stalled peptide with the appearance of a ‘smear’ of CAT tailed peptides. Interestingly, combined deletion of both *JLP2* and *LTN1* resulted in a further increase in CAT tailing and accumulation of high molecular weight CAT tail-dependent aggregates migrating close to the well during SDS-PAGE^44,53–56^ (Fig. 1c). Both CAT tails and aggregates were lost with deletion of *RQC2* (Fig. 1c), confirming aggregate formation was CAT tail dependent. Additionally, TEV protease cleavage collapsed the smear into a single discrete band, further supporting the interpretation of the smear as CAT tailed species (Supplementary Fig. S1b). To assess whether this observation was generalizable to conditions that cause ribosome stalling, we also used a GFP-TEV-Rz reporter (pGTRz) containing a self-cleaving hammerhead ribozyme sequence that results in a truncated mRNA without a stop codon, causing ribosomes to stall at the very 3’ end^21^. Although CAT tail ‘smears’ were not readily visible for this reporter, we consistently observed similar increases in CAT tail-dependent aggregate formation associated with loss of *JLP2* in the *ltn1Δ* background (Supplementary Fig. S1c), suggesting that Jlp2 generally counteracts CAT tailing in the absence of Ltn1. Since the visibility of CAT tailed smears varied between reporters, we used CAT tail-dependent aggregate formation as a more robust and consistent readout for changes in CAT tailing in subsequent experiments. We also noted a decrease in the level of the unaggregated reporter protein in the *jlp2Δltn1Δ* double deletion strain (Supplementary Fig. S1c) which may be due to the combined effect of increased reporter aggregation (Supplementary Fig. S1c) and a decrease in reporter mRNA level (Supplementary Fig. S1d). Collectively, these results show that loss of *JLP2* results in an increased accumulation of CAT tail-dependent aggregates.

To determine whether the observed increase in CAT tail-dependent aggregates in *jlp2Δltn1Δ* was caused by the absence of Ltn1-dependent nascent peptide ubiquitination rather than an indirect effect of *LTN1* deletion, we employed a lysine-free stalling reporter devoid of lysines available for Ltn1-mediated ubiquitination (GFP_Lys-free_-3xHA-TEV-R12-RFP) and a matching control with lysine-containing GFP (GFP_Lys_-3xHA-TEV-R12-RFP)^37^. For the lysine-free reporter, any effect on CAT tailing is directly attributable to absence of Jlp2 without perturbing *LTN1*. Indeed, for the lysine-free, but not the lysine-containing reporter, deletion of *JLP2* was sufficient to drive accumulation of CAT tail-dependent aggregates, fully suppressed by plasmid-based Jlp2 overexpression, which even eliminated the (mild) accumulation of CAT tailed products in the *ltn1Δ* single knockout strain (Fig. 1d). Furthermore, overexpression of Jlp2 suppressed CAT tail-dependent aggregates in a strain expressing catalytically inactive Ltn1^31^ (Supplementary Fig. S1e). Together, these findings show that Jlp2 counteracts excessive CAT tailing and accumulation of CAT tail-dependent aggregates of RQC substrates that escape Ltn1-mediated ubiquitination.

Finally, to define the extent to which Jlp2 counteracts CAT tailing, we used a stalling reporter in which lysines are encoded only at the C terminus and remain buried within the ribosomal exit tunnel upon ribosome stalling, inaccessible to Ltn1 (GFP_Lys-free_-3xHA-TEV-(A)30-Rz^37^). For Ltn1-dependent ubiquitination and proteasomal degradation, CAT tailing is required to expose those lysines^37^ (see Supplementary Fig. S1f for schematic). Accordingly, if Jlp2 blocked CAT tailing completely, the reporter protein would be stabilized to levels seen in *ltn1Δ* even in the presence of Ltn1^37^. Quantification after treatment with TEV protease (to remove poly(A)-encoded lysines and CAT tails) revealed that overexpression of Jlp2 in wild-type cells did not stabilize the reporter protein (Supplementary Fig. S1f) indicating that CAT tailing is modulated rather than fully abolished. Collectively, these results suggest that Jlp2 modulates the extent of CAT tailing under conditions in which Ltn1-dependent ubiquitination is compromised without interfering with canonical RQC complex function.

### Jlp2 is required for peptide release from aberrant peptidyl-tRNA in the absence of RQC complex proteins

Mechanistically, we first examined whether Jlp2 regulates CAT tailing by modulating either Rqc2 levels or the 60S:peptidyl-tRNA complex binding of Rqc2. However, overexpression of Jlp2 in a *jlp2Δltn1Δ* strain neither reduced Rqc2 levels (Supplementary Fig. S2a); nor displaced Rqc2 from the 60S fraction in sucrose gradients, either in absence (Supplementary Fig. S2b) or presence (Supplementary Fig. S2c) of CHX in the *ltn1Δ* background. Instead, unlike Rqc2, Jlp2 associated most prominently with the 80S fraction, further supporting the observation that Jlp2 does not compete with Rqc2 to counteract CAT tailing. This observation is consistent with the conclusion that Jlp2 does not interfere with RQC complex function. 60S-associated aberrant peptidyl-tRNA act as substrates for CAT tailing^30,36^. Vms1 (ANKZF1 in mammals) functions as a release factor for ubiquitinated nascent peptides tethered to the 60S via their tRNAs^28,41–43^. Deleting *LTN1* diminished Vms1’s 60S association^41^, indicating nascent peptide ubiquitination may recruit Vms1 to the 60S. Consistent with this, CHX hypersensitivity of *vms1Δ* is abolished by additional deletion of *LTN1* and other RQC complex factors^42^. In contrast, we found that deletion of RQC components exacerbates the CHX hypersensitivity associated with loss of Jlp2 (Fig. 1b and Supplementary Fig. S1a). Nevertheless, both Jlp2 (Fig. 1c and d) and Vms1^44^ counteract CAT tailing. Therefore, we hypothesized that unlike Vms1, Jlp2 may be an RQC complex-independent release factor, representing an alternative quality control mechanism operating in the absence of K48-linked nascent peptide polyubiquitination.

To test this hypothesis, we examined strains lacking *JLP2* and/or RQC complex factors for accumulation of tRNA-conjugated species of the stalled reporter protein using Western blotting under neutral pH conditions, which preserve the peptidyl-tRNA ester bond otherwise hydrolyzed during standard SDS-PAGE. We employed a Protein-A-NonStop (PrANS) reporter^57^ that has previously been used to visualize peptidyl-tRNA bands with high clarity in the context of RQC factor deletants^34,41^. Consistent with our hypothesis, deletion of *JLP2* resulted in an increase in the peptidyl-tRNA band in the background of strains lacking various RQC complex members (Fig. 2a). RNase treatment abolished this band, with a concomitant increase of the free peptide (Supplementary Fig. S2e), confirming its identity as a peptidyl-tRNA species.

**Figure 2.**
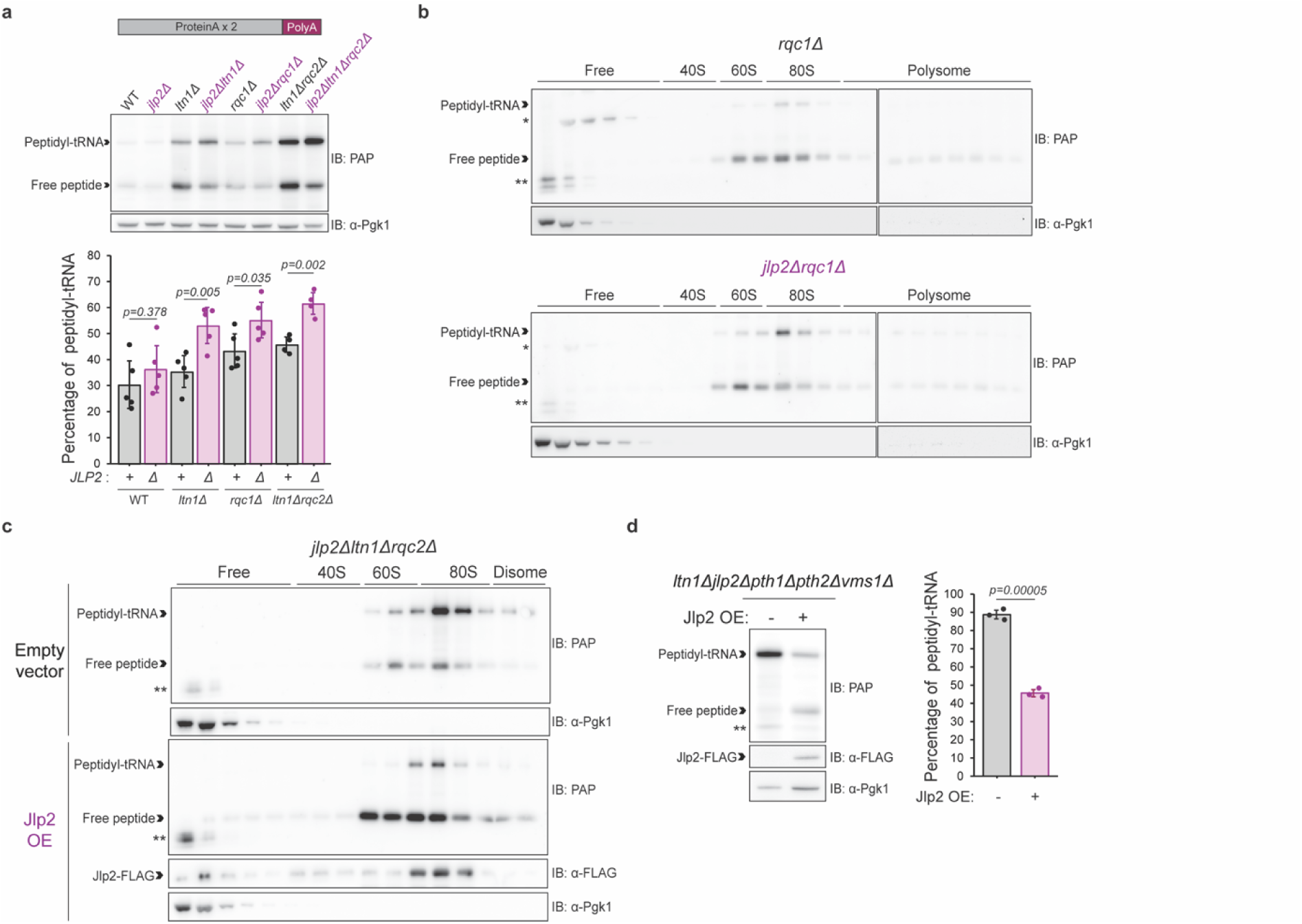
Jlp2 is required for the release of peptidyl-tRNA of ribosome stalling substrates in the absence of canonical RQC. a. Schematic representation of the PrANS reporter and Western blot analysis of peptidyl-tRNA species of PrANS reporter in WT, *jlp2Δ, ltn1Δ, jlp2Δltn1Δ, rqc1Δ, jlp2Δrqc1Δ, ltn1Δrqc2Δ* and *jlp2Δltn1Δrqc2Δ* and quantification of percentage of PrANS peptidyl-tRNA relative to total PrANS (free peptide + peptidyl-tRNA). Student’s t-test (two-tailed, heteroscedastic) was used to calculate the p-value. N=3 biological replicates, n=1 or 2 technical replicates. b. Sucrose density gradient fractionation and Western blot analysis of peptidyl-tRNA and free peptide species of PrANS reporter in *rqc1Δ* and *jlp2Δrqc1Δ.* The ∼40 kDa band migrating in the free fraction, denoted by asterisk (*) denotes a non-reporter specific (Supplementary Fig. S2d), RNase-resistant (Supplementary Fig. S2h) band that is inconsistently observed across experiments. Double-asterisk (**) denotes unassigned, reporter-specific signals (Supplementary Fig. S2d). c. Overexpression of Jlp2 counteracts the accumulation of peptidyl-tRNA of RQC substrates in the absence of RQC complex activity. Sucrose density gradient fractionation and Western blot analysis of peptidyl-tRNA and free peptide species of PrANS reporter in *jlp2Δltn1Δrqc2Δ,* containing either an empty vector or a Jlp2-FLAG overexpression (OE) vector. Double-asterisk (**) denotes unassigned reporter-specific signals (Supplementary Fig. S2d). d. Western blot analysis of peptidyl-tRNA and free peptide species of PrANS reporter in *ltn1Δjlp2Δpth1Δpth2Δvms1Δ* containing either an empty vector or Jlp2-FLAG OE vector and quantification of percentage of PrANS peptidyl-tRNA relative to total PrANS (free peptide + peptidyl-tRNA). Student’s t-test (two-tailed, heteroscedastic) was used to calculate the p-value. N=3 biological replicates.

We also examined peptidyl-tRNA species using the pGTRz reporter. Here, a faint, RNase-sensitive peptidyl-tRNA band appeared only upon deletion of *JLP2* in addition to *LTN1* or *RQC1,* with no visible peptidyl-tRNA band in the single knockout strains (Supplementary Fig. S2f and g). This is consistent with observations of others, where GFP-based reporters showed faint peptidyl-tRNA bands (if any at all)^34,58,59^ on Western blots. Together, these results show that absence of Jlp2 results in the stabilization of aberrant peptidyl-tRNA in the absence of RQC complex activity.

To examine whether the PrANS peptidyl-tRNA was still tethered to the ribosome, we performed sucrose density gradient fractionation in *rqc1Δ* or *rqc1Δjlp2Δ* cells. Peptidyl-tRNA was found in the ribosomal fractions with most of the peptidyl-tRNA species in the 80S (monosome) fraction, with the deletion of *JLP2* showing increased stabilization of peptidyl-tRNA species (Fig. 2b). Indeed, 80S associated-nascent peptides in RQC factor deletants have been observed consistently in literature and attributed to spontaneous subunit rejoining^25,31,41,56^. Further, overexpression of Jlp2 resulted in the release of the PrANS peptide from tRNA in cells lacking endogenously expressed Jlp2, Ltn1 and Rqc2 (Fig. 2c). Again, the majority of both peptidyl-tRNA and Jlp2 were associated with the 80S fraction (Fig. 2c). Despite efficient conversion of peptidyl-tRNA to free peptide, we did not observe active drop-off of the free peptide to lower density fractions. Instead, we observed weak bands for free PrANS peptide in the lower fractions (Fig. 2c), in line with the expectation that active extraction of the stalled peptide by the Cdc48 complex does not occur in the absence of nascent peptide ubiquitination. Taken together, these results suggest that Jlp2 may be a release factor that acts on aberrant peptidyl-tRNA in the absence of RQC complex activity, including both, nascent peptide ubiquitination and CAT-tailing.

To determine whether the 80S-associated substrates indeed represented spontaneously rejoined subunits formed after initial ribosome splitting and not translating/stalled ribosomes on mRNA, we examined Jlp2-dependent peptide release in cells lacking Hel2 (prohibiting ribosome splitting by the RQT complex and subsequent RQC)^9,17,60^ and Dom34 (prohibiting alternate pathways, such as splitting ribosomes stalled at the 3’-end of mRNA or empty ribosomes)^18,26,61^. Indeed, we found that Jlp2 overexpression in a *jlp2Δhel2Δdom34Δ* strain did not result in the conversion of PrANS peptidyl-tRNA to free peptide compared to the empty vector control (Supplementary Fig. S2i). These results indicate that Jlp2-dependent peptide release predominantly occurs downstream of RQT and/or Dom34-dependent ribosome splitting and support the conclusion that the peptidyl-tRNA:80S substrates of Jlp2 indeed represent spontaneously rejoined subunits and not translating/stalled ribosomes on an mRNA.

### Jlp2 functions independently of release factors Vms1, Pth1 and Pth2

Next, we interrogated potential interactors of Jlp2. Jlp2-FLAG expressed from a plasmid in a *jlp2Δ* strain co-immunoprecipitated proteins of the large and small ribosomal subunits (Supplementary Fig. S2j, see Supplementary Table S1), among the most abundant proteins in the analysis. In line with the RQC complex-independent function of Jlp2, RQC factors Ltn1, Rqc1, Rqc2 and Cdc48 complex proteins were not enriched in the Jlp2-FLAG pulldown (Supplementary Fig. S2j). Among the interactors was peptidyl-tRNA hydrolase Pth1, but not Pth2 (Supplementary Fig. S2j), whereas other mitochondrial peptidyl-tRNA hydrolases Rso55 (Pth3) and Pth4 were not quantifiable, hence unlikely to interact significantly. Bacterial Pth^62^ and mammalian PTRH1^28^ have been shown to release nascent peptides arising from ribosome stalling, potentially suggesting a functional link between Pth1 and Jlp2. Recently, yeast Pth2 has also been proposed to act as a release factor for stalled products of mitochondrial translocation-coupled translation^58^ and therefore still remained a candidate for functional overlap. Jlp2 also pulled down Vms1, but not its cofactor Arb1^43^(Supplementary Fig. S2j).

To determine whether Jlp2-dependent peptide release required these known release factors, we examined peptidyl-tRNA accumulation in *ltn1Δ* cells lacking *JLP2, PTH1, PTH2* and *VMS1* in all possible combinations (Supplementary Fig. S2k). Individual deletions of *JLP2, VMS1* or *PTH2* but not *PTH1* resulted in increased stabilization of PrANS peptidyl-tRNA, with their combined deletions (again with the exception of *PTH1*) producing additive effects on peptidyl-tRNA stability (Supplementary Fig. S2k), indicative of independent functioning of Jlp2, Vms1 and Pth2 and no observable role for Pth1 in the quality control of cytoplasmic substrates. Notably, compared to the combined deletion of *LTN1, PTH1, PTH2* and *VMS1*, the additional deletion of *JLP2* resulted in a near-complete stabilization of peptidyl-tRNA (Supplementary Fig. S2k), supporting the notion that Jlp2 functions independently of all these factors. Consistent with this interpretation, overexpression of Jlp2 in an *ltn1Δjlp2Δpth1Δpth2Δvms1Δ* strain was sufficient to drive the conversion of peptidyl-tRNA to free peptide (Fig. 2d), confirming that Jlp2-dependent peptide release does not require these factors. Additionally, the top interacting partners of Jlp2 identified by mass spectrometry - Pro1 and its uncharacterized paralog Yhr033w, were dispensable for Jlp2-mediated peptide release (Supplementary Fig. S2l). Collectively, these results strongly suggest Jlp2 as a novel release factor with its own catalytic residues and/or architecture.

## A conserved histidine is required for Jlp2-mediated peptide release

Since Jlp2 did not present any structural resemblance to or motifs found in known nucleases or hydrolases, we examined its evolutionary sequence conservation from yeast to animals and plants (Supplementary Fig. S3a), as well as structural features (Supplementary Fig. S3b-d) to gain deeper insights into Jlp2’s peptide release mechanism. We used the AlphaFold-predicted structure of Jlp2 (AF-P40206-F1-v6)^52^, that closely matches the x-ray structure of its mammalian homolog CCDC25 (PDB: 7EOE, Chain A) (Supplementary Fig. S3b), particularly within the highly conserved regions including NFACT-R domain (Supplementary Fig. S3c and d), supporting the model’s reliability. The C-terminal region diverged in orientation beginning at yeast residue 157, coinciding with a predicted flexible segment, but retained an overall similar fold thereafter (Supplementary Fig. S3b and d).

Mapping sequence conservation of Jlp2 onto the structure revealed a prominent cavity within the NFACT-R domain, lined by highly conserved residues (Fig. 3a and Supplementary Fig. S3e), similarly present in CCDC25 (Supplementary Fig. S3f). To assess its functional relevance, we introduced alanine substitutions at conserved residues lining this cavity, with the exception of D46 which was mutated to asparagine. Additionally, several highly conserved residues outside of this cavity were also mutated to alanine. We then evaluated peptide release upon overexpression of mutant or WT Jlp2 in the *ltn1Δjlp2Δpth1Δpth2Δvms1Δ* strain. Although substitutions at several highly conserved positions altered peptide release efficiencies, only substitution of histidine-44 (H44) completely abolished conversion of peptidyl-tRNA to free peptide (Fig. 3b). Evaluation of selected mutants by sucrose density gradient fractionation in *jlp2Δltn1Δrqc2Δ* yielded consistent results, with only the H44A substitution showing a loss of Jlp2-mediated peptide release (Supplementary Fig. S3g). Importantly, the H44A mutant retained association with the 80S ribosomal fraction, indicating that loss of activity is not attributable to impaired ribosome binding (Supplementary Fig. S3g). As the Jlp2 H44A mutant and its orthologous mutation in CCDC25 were predicted to exhibit a modest reduction in folding stability (Extended Data Table 2), we introduced alternative substitutions at position 44 (H44F and H44Y) and two neighbouring residues (H52F/Y and N27I/L), all of which were predicted to have negligible effects on (or even improve) folding stability (Supplementary Table S2). Only substitutions at position 44 abolished Jlp2-dependent peptide release, whereas substitutions at nearby positions did not (Fig. 3c). Importantly, the H44F mutant also retained ribosome binding similar to WT Jlp2 (Fig. 3d). Together, these results show that H44 is essential for Jlp2-dependent peptide release.

**Figure 3.**
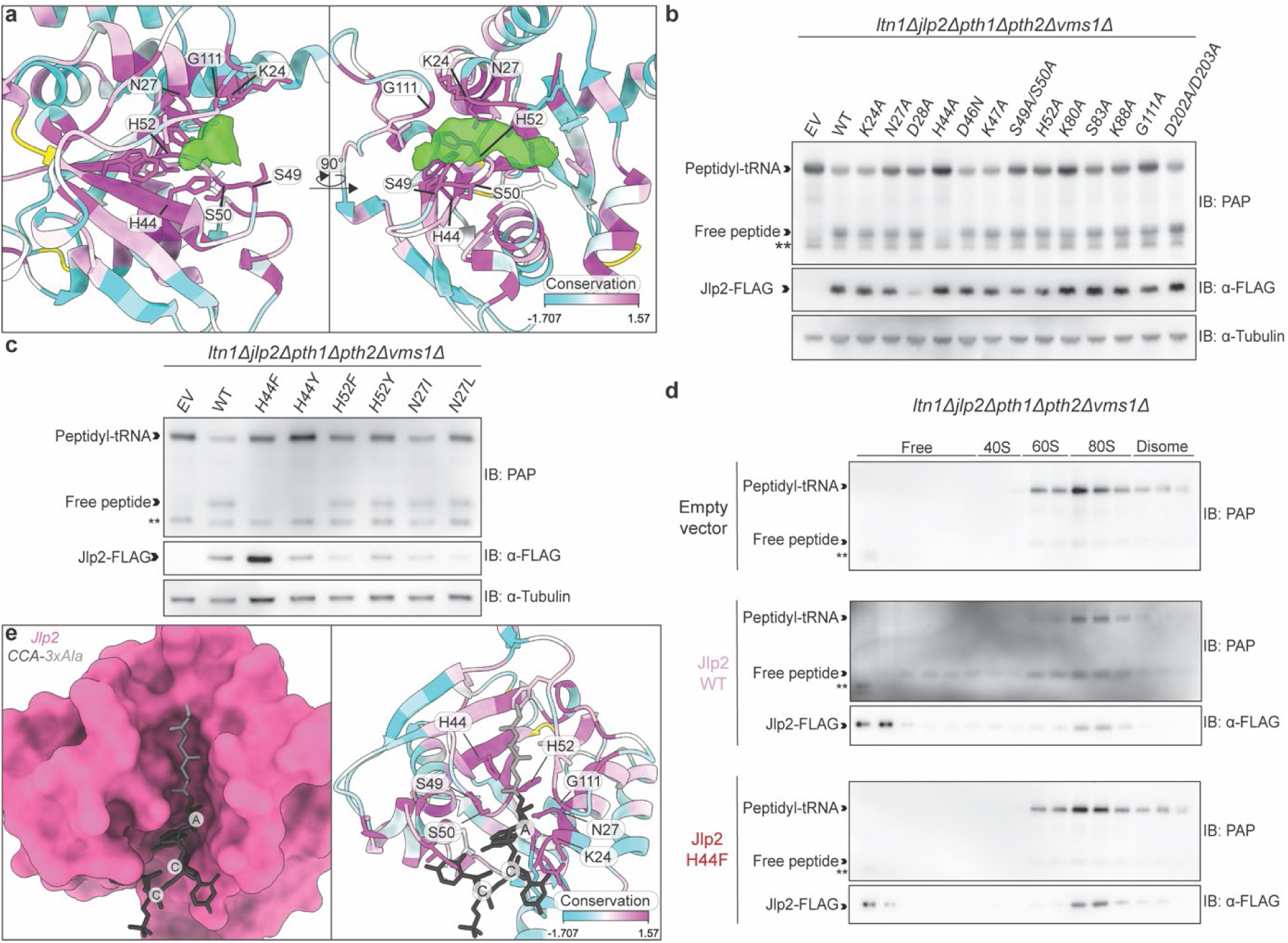
Identification of a conserved residue essential for the release factor activity of Jlp2 through sequence and structure conservation analysis. a. AlphaFold-predicted structure of Jlp2 [AF-P40206-F1-v6 (average pLDDT= 94.69)]^52^ colored as per sequence conservation (see scale) highlighting a prominent cavity (green density) lined with highly evolutionarily conserved residues. Residues shown in yellow correspond to positions for which sequence conservation values could not be determined. b. H44 is essential for Jlp2-mediated peptide release from tRNA. Western blot analysis of peptidyl-tRNA and free PrANS reporter species in *ltn1Δjlp2Δpth1Δpth2Δvms1Δ* containing either an empty vector or a plasmid expressing WT Jlp2 or the indicated Jlp2 substitution mutants. c. Alternative substitutions at position 44 but not at neighbouring positions abolish Jlp2-mediated peptide release. Western blot analysis of PrANS peptidyl-tRNA and free PrANS reporter species in *ltn1Δjlp2Δpth1Δpth2Δvms1Δ* containing an empty vector or a plasmid expressing WT mutants or the indicated substitution mutants of Jlp2. d. The catalytically inactive H44F mutant retains association with the 80S ribosomal fraction. Sucrose density gradient fractionation and Western blot analysis of PrANS peptidyl-tRNA and free PrANS reporter species and ribosomal association of Jlp2 H44F mutant in an *ltn1Δjlp2Δpth1Δpth2Δvms1Δ* strain. e. OpenFold3-predicted model of Jlp2 associated with CCA-3xAla ligand (Model 1). Left: Surface representation of Jlp2 (pink) with CCA (black) - 3xAlanine (grey) ligand, right: Ribbon representation of Jlp2 (colored according to sequence conservation as in panel a) in association with CCA-3xAla (grey-black). Residues at the predicted interface are labelled. Model confidence metrics: IPTM = 0.856047, PTM = 0.885398, average PLDDT = 90.754242, GPDE = 0.671759 and Sample ranking score = 0.895571 (see Supplementary Fig. S3h and i for per residue/atom pLDDT scores).

Next, we used the AlphaFold3^64^ reproduction-OpenFold3^65,66^ to perform a predictive modelling of the interaction between Jlp2 and a minimal peptidyl-tRNA ligand representing the 3′-CCA end of tRNA linked to three alanine residues by an ester bond at the 3’-Oxygen (CCA-3×Ala). OpenFold3 modelled the ligand within the conserved cavity of the NFACT-R domain (Fig. 3a) of Jlp2 as the predictedinteraction interface with the peptidyl-tRNA ester bond positioned in close proximity to H44 (Fig. 3e, Supplementary Fig. S3h-k).

Collectively, our mutagenesis data identify H44 as an essential residue for Jlp2-dependent peptide release, supported by structural modelling suggesting a mechanism in which the conserved NFACT-R cavity mediates peptidyl-tRNA engagement and H44 as the catalytic residue that facilitates peptidyl-tRNA hydrolysis.

### Jlp2 directly catalyzes peptide release from ribosome-associated aberrant peptidyl-tRNA

To determine whether Jlp2 directly catalyzes peptide release on peptidyl-tRNA:ribosome complexes, we established an on-bead *in vitro* reconstitution system (see Methods). We reconstituted PrANS peptidyl-tRNA:ribosome complexes enriched from *jlp2Δltn1Δrqc2Δ* strains under two conditions: a “high salt” condition to deplete weakly associated proteins or a “physiological salt” condition, intended to preserve associated factors (Fig. 4a, see Methods for details) with purified Jlp2. *In vitro* reconstitution reactions demonstrated that WT but not H44A Jlp2 catalyzed conversion of peptidyl-tRNA to free peptide (Fig. 4b) requiring neither Ltn1-dependent ubiquitination nor Rqc2-mediated CAT tailing. The absence of any apparent differences between “high salt” and “physiological salt”-enriched complexes (Fig. 4b) further supports the notion that Jlp2’s activity is not dependent on accessory factors co-purifying with the peptidyl-tRNA:ribosome complexes. We also examined Jlp2-mediated peptide release using high salt-washed PrANS peptidyl-tRNA:ribosome complexes from *ltn1Δjlp2Δpth1Δpth2Δvms1Δ* strains. Consistent with our *in vivo* results, only WT but not H44F Jlp2 facilitated peptide release (Fig. 4c), confirming that H44 is essential for Jlp2-mediated peptide release. To assess whether Jlp2 is capable of peptide release from both 60S- and 80S-associated peptidyl-tRNA in the absence of other release factors, we performed sucrose density gradient fractionation to separately enrich for 60S and 80S complexes followed by enrichment of high salt-washed PrANS peptidyl-tRNA:ribosome complexes from *ltn1Δjlp2Δpth1Δpth2Δvms1Δ* strains for on-bead *in vitro* reconstitution (see Methods). Indeed, Jlp2 catalyzed peptide release on both 60S and 80S complexes with comparable efficiencies (Fig. 4d). Analogous experiments using purified Jlp2-FLAG with either peptidyl-tRNA:60S or peptidyl-tRNA:80S complexes from *jlp2Δltn1Δrqc2Δ* enriched using only sucrose density gradient fractionation under “high salt” and “physiological salt” conditions yielded similar results (Supplementary Fig. S4a and b). Across all *in vitro* and *in vivo* experiments, a fraction of peptidyl-tRNA remained unreleased, and prolonged incubation of peptidyl-tRNA:80S complexes with Jlp2 did not facilitate further release (Supplementary Fig. S4c), suggesting that a subpopulation of peptidyl-tRNA:ribosome complexes may adopt a Jlp2-resistant state or conformation.

**Figure 4.**
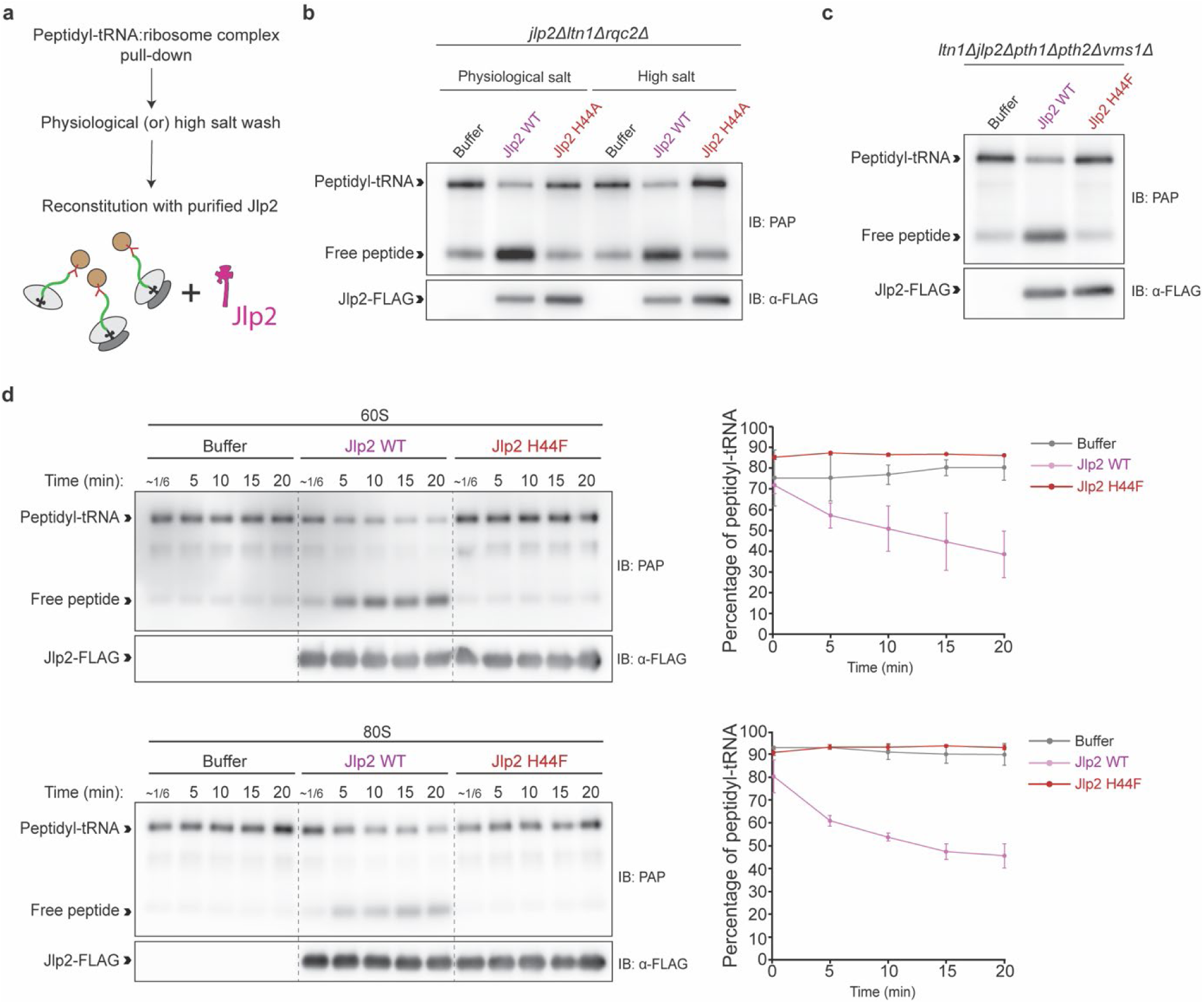
Jlp2 directly catalyzes peptide release from ribosome-associated aberrant peptidyl-tRNA. a. Schematic overview of the IgG magnetic bead-based enrichment strategy for PrANS peptidyl-tRNA:ribosome complex used for on-bead *in vitro* reconstitution with purified Jlp2. Peptidyl-tRNA ribosomes were pulled down using IgG (red) magnetic beads (brown) and washed under high salt or physiological salt conditions and reconstituted with purified Jlp2. b. Jlp2 catalyzes peptide release from ribosome-associated peptidyl-tRNA *in vitro* in the absence of both Ltn1-mediated ubiquitination and Rqc2-mediated CAT tailing. Western blot analysis of on-bead *in vitro* reconstitution reactions using purified Jlp2 or the H44A mutant and peptidyl-tRNA:ribosome complexes enriched from *jlp2Δltn1Δrqc2Δ* cells under high salt or physiological salt conditions. c. Jlp2-mediated peptide release is independent of Vms1, Pth1 and Pth2. Western blot analysis of on-bead *in vitro* reconstitution reactions using purified Jlp2 and peptidyl-tRNA:ribosome complexes enriched under high salt conditions from *ltn1Δjlp2Δpth1Δpth2Δvms1Δ* cells. d. Jlp2 catalyzes peptide release from both 60S and 80S-associated peptidyl-tRNA with comparable efficiency. Time course Western blot analysis of on-bead *in vitro* reconstitution reactions using Jlp2 and peptidyl-tRNA:60S or 80S complexes both purified from *ltn1Δjlp2Δpth1Δpth2Δvms1Δ* cells for peptidyl-tRNA release and quantification of percentage of peptidyl-tRNA. Ribosomal complexes were fractionated using sucrose density gradients followed by IgG magnetic bead-based enrichment under high salt conditions (n=2 independent experiments). All samples were resolved on the same gel and the dotted lines indicate boundaries between conditions for clarity.

To verify whether our fractionated peptidyl-tRNA:80S remained intact during the experiment, we performed analytical sucrose density gradient fractionation after the *in vitro* reactions, under buffer conditions matching those of the *in vitro* reconstitution reactions. Absorbance profiles showed that the isolated 80S fractions contained a minor contaminating 60S but no 40S absorbance peaks, suggesting that those were 60S already present in the initial purification, and not resulting from splitting of 80S into 60S and 40S (Supplementary Fig. S4d). Nevertheless, nearly all peptidyl-tRNA remained in the 80S fraction (Supplementary Fig. S4d, bottom), and peptidyl-tRNA conversion to free peptide occurred in both fractions demonstrating that Jlp2 can act on peptidyl-tRNA:80S complexes.

Taken together, these results establish Jlp2 as a *bona fide* release factor capable of directly catalyzing peptide release from aberrant peptidyl-tRNA.

### Release factor activity of Jlp2 is required for protection against translation elongation stress

To determine whether Jlp2’s release factor activity is required to suppress CAT tailing, we compared WT Jlp2 to the catalytically inactive H44A mutant for their ability to suppress CAT tail-dependent aggregate formation in the *jlp2Δltn1Δ* background. Indeed, loss of release factor activity abolished the suppression of CAT tail-dependent aggregation (Fig. 5a), demonstrating that Jlp2’s release factor activity is required to counteract CAT tailing.

**Figure 5.**
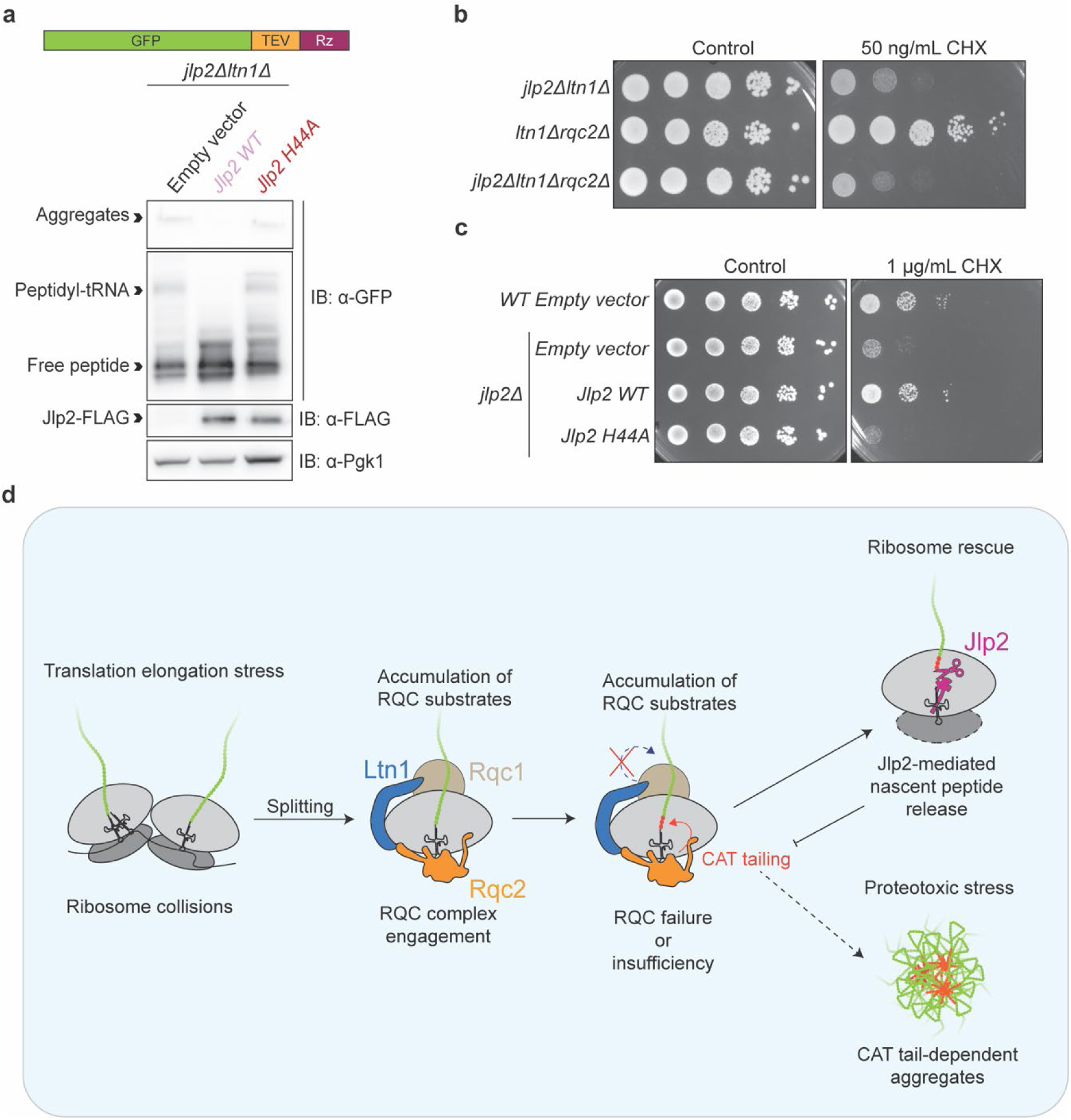
The release factor activity of Jlp2 is required for its role in translation quality control. a. Release factor activity of Jlp2 is required for the suppression of CAT tail-dependent aggregate formation. Western blot analysis of the pGTRz reporter for peptidyl-tRNA species and aggregate accumulation in *jlp2Δltn1Δ* cells containing either an empty vector or overexpression vectors for wild-type or H44A mutant Jlp2. b. CHX hypersensitivity in the absence of *JLP2* and *LTN1* is not attributable to excessive CAT tailing and CAT tail-dependent aggregate formation. Growth of *jlp2Δltn1Δ, ltn1Δrqc2Δ* and *jlp2Δltn1Δrqc2Δ* spotted in serial dilutions on CHX-containing plates, imaged after 3 days. c. The release factor activity of Jlp2 is required for protection against translation elongation stress. Spot assay showing the growth of WT cells containing an empty vector and *jlp2Δ* cells containing either an empty vector, or overexpression vectors for WT or H44A Jlp2 spotted in serial dilutions on CHX-containing plates, imaged after 4 days and 8 days for control and 1 µg/mL respectively. d. Model depicting the role of Jlp2 in TQC. Translation elongation stress induces ribosome collisions, which trigger ribosome splitting and generate peptidyl-tRNA:60S complexes that are normally processed by the RQC complex. Such complexes may also spontaneously reassociate with the 40S subunit to form 80S monosomes. Upon failure or insufficiency of RQC complex activity, Jlp2 rescues ribosomal subunits by catalyzing peptide release from both peptidyl-tRNA:60S and spontaneously reassociated peptidyl-tRNA:80S complexes, thereby restricting excessive CAT tail extension and preventing the formation of CAT tail-dependent protein aggregates.

Having identified two distinct but related functions of Jlp2 - suppression of CAT tail-dependent aggregation and peptide release from aberrant ribosome-associated peptidyl-tRNA – we now asked which of both functions protects against CHX-induced translation elongation stress. Additional deletion of *RQC2* did not suppress the CHX-hypersensitivity of the *jlp2Δltn1Δ* strain (Fig. 5b), suggesting that excessive CAT tail-mediated aggregation of RQC substrates is not the primary determinant of CHX induced translation elongation stress (Fig. 1b). On the other hand, only overexpression of WT Jlp2, but neither the H44A mutant (Fig. 5c) nor the H44F mutant (Supplementary Fig. S5a) restored growth of *jlp2Δ* strains in presence of CHX, suggesting that the specific requirement of Jlp2-dependent peptide release may serve an additional purpose beyond regulating CAT tailing and propose that the rescue of ribosomal subunits from aberrant peptidyl-tRNA occupancy through the release factor activity of Jlp2 constitutes the principal mechanism by which Jlp2 protects against translation elongation stress.

Collectively, our results define Jlp2 as a novel TQC release factor that protects cells against ribosome stalling stress by catalyzing the release of aberrant nascent peptides from ribosome-bound tRNA, thereby potentially rescuing stalled ribosomal subunits and restricting excessive CAT tailing to prevent the aggregation of aberrant nascent peptides (Fig. 5d).

## Discussion

Ribosome stalling and consequent aberrant peptide accumulation challenge cellular proteostasis. Depending on multi-factor coordination, canonical RQC is vulnerable to failure. Here, we identify and characterize Jlp2 as a TQC factor operating as a fail-safe for RQC substrates escaping canonical processing. We demonstrate that Jlp2 catalyzes nascent peptide release from ribosome-associated tRNA independently of the RQC complex and previously characterized TQC release factors Vms1, Pth1 and Pth2, rescuing blocked ribosomal subunits and restricting CAT tail-dependent aggregate formation. Instead of interfering with RQC complex activity, Jlp2 acts downstream or in parallel, becoming critical when K48-linked nascent peptide ubiquitination is compromised (Fig. 1b, 1d, 2a and 2b). This situation arises in physiologically relevant scenarios, including reduced lysine accessibility due to nascent peptide composition, -folding or co-translational translocation; or RQC machinery mutation, scarcity or saturation upon translation stress. The finding that deletion of *JLP2* alone confers translation inhibitor hypersensitivity even with a functional RQC machinery (Fig. 1b and Supplementary Fig. S1a) underscores Jlp2’s importance as a constitutive safeguard rather than a purely compensatory factor.

Without Ltn1-mediated ubiquitination, CAT tailing can trigger peptide degradation. In yeast, this requires Hul5-mediated ubiquitination^38^. In mammals, CAT tails are recognized by C-end rule ubiquitin ligases, including Pirh2 and the CRL2-KLHDC10 complex, promoting proteasomal degradation^39,67^. However, excessively extended CAT tails mediate proteasome-resistant aggregation^40,56^. Protein aggregation contributes to neurodegenerative disease^68–70^ and loss of LISTERIN causes neurodegenerative phenotypes, rescued by depletion of NEMF, indicating excessive CAT tailing as a pathogenic driver^71,72^. In this context, our finding that Jlp2 limits CAT tailing (Fig. 1d) without completely abolishing it (Supplementary Fig. S1f) is noteworthy. Increased CAT tail lengths are associated with the propensity to form CAT tail-dependent aggregates whereas shorter CAT tails facilitate degradation^38,56^, correlating with the observed increase in CAT tail lengths and aggregates in our experiments (Fig 1c and d). By releasing the nascent peptide from the ribosome before CAT tails reach pathological lengths, Jlp2 may preserve the degron function of CAT tails, while preventing aggregation. Jlp2 achieves this without competing with Rqc2 for 60S binding (Supplementary Fig. S2b, c) or affecting canonical RQC complex function (Supplementary Fig. S1f), suggesting a substrate-level rather than complex-level mode of regulation.

Although Jlp2 counteracts excessive CAT tailing and aggregation (Fig. 1c, d), abolishing CAT tailing did not rescue the CHX hypersensitivity of *jlp2Δltn1Δ* cells (Fig. 5b). Overexpression of WT, but not catalytically inactive Jlp2 H44A and H44F rescued CHX sensitivity in *jlp2Δ* strains (Fig. 5c and Supplementary Fig. S5a), directly linking the protective function of Jlp2 to its peptidyl-tRNA release activity. These observations suggest that ribosomal subunit rescue from aberrant peptidyl-tRNA occupancy is the principal mechanism by which Jlp2 confers protection against translation elongation stress. In addition, by releasing nascent peptides from tRNA, Jlp2 may also facilitate their degradation, including through CAT tail-mediated degradation without Ltn1-catalyzed ubiquitination^38,39,56,67^, contributing to proteostasis through a secondary route.

Vms1 regulates CAT tailing of stalled mitochondrial substrates^43,44^, with its efficacy as a release factor hinted to depend on Ltn1-mediated RQC substrate ubiquitination^28,41,42^. Additionally, mammalian PTRH1 - ortholog of yeast Pth1 - has been shown to release non-ubiquitinated RQC substrates *in vitro*^28^, while Pth releases bacterial RQC substrates^62^. Since yeast Pth1 resides in the mitochondrial matrix^58,73^ and mammalian PTRH1 also localized to the mitochondrion^74^, their activities on cytoplasmic substrates *in vivo* remain unresolved. Pth2 is associated with the outer mitochondrial membrane, facing the cytosol and proposed to act as a release factor of mitochondrially targeted RQC substrates without affecting CAT tailing^58^. Our experiments show that Pth2, but not Pth1, also acts as a release factor on cytosolic substrates (Supplementary Fig. S2k), and that Jlp2, Pth2 and Vms1 may be the only release factors acting on cytoplasmic stalling substrates, their combined absence resulting in near-complete stabilization of reporter peptidyl-tRNA (Supplementary Fig. S2k). Residual conversion to free peptide in *ltn1Δjlp2Δpth2Δvms1Δ* likely reflects spontaneous peptidyl-tRNA hydrolysis due to experimental factors such as heat and/or pH. Our work suggests that these factors may be redundant to some extent (Supplementary Fig. S2k), while the remaining peptidyl-tRNA in our *in vitro* reconstitution reactions (Fig. 4) suggests a defined substrate scope for Jlp2 - and possibly the other release factors as well. With our definition of all cytoplasmic, noncanonical release factors, it will be interesting to further characterize their substrate specificities and redundancies.

The complete loss of Jlp2’s activity upon H44 mutation (Fig. 3b and c), combined with H44’s predicted proximity to the peptidyl-tRNA ester bond (Fig. 3e) is consistent with a catalytic role. H44 being the essential catalytic residue (Fig. 3) places Jlp2 in a class distinct from canonical eukaryotic peptide release factors. During canonical translation termination, eukaryotic release factor eRF1 and its bacterial ortholog employ a conserved GGQ motif for peptidyl-tRNA ester bond hydrolysis^75–79^. Recently, cryo-EM structures of (bacterial) RF1 in *Thermus thermophilus* with peptidyl-tRNA^75^ showed that the NH (of the amide bond) of the GGQ-motif’s glutamine forms a hydrogen bond with the carbonyl-oxygen, causing the CCA-adenosine’s ribose to be in a C3’-endo pucker, suggested to facilitate a nucleophilic attack of the 2’-OH on the carbonyl carbon^75^. Our co-folding predictions for Jlp2 employing a CCA-3xAla peptidyl-tRNA mimic suggest a hydrogen bond between H44’s ring *N^τ^*H and the carbonyl-oxygen of the ester bond, with the terminal adenosine-ribose in a C3’-endo pucker (Supplementary Fig. S3k), suggesting a plausible mechanism for Jlp2’s histidine-mediated catalysis. Of note, Vms1 is proposed to employ a GGQ-related GSQ motif (ANKZF1 possessing an orthologous TAQ motif), but instead of hydrolysing the peptidyl-tRNA ester bond, Vms1 endonucleolytically cleaves tRNA near the CCA-end^28,43,80^. Although a similar mechanism cannot be excluded for Jlp2, the co-folding predictions do not support this possibility. To the contrary, Jlp2’s dependence on histidine H44 is analogous to Pth, where a histidine, as a general base, activates water for nucleophilic attack on the peptidyl-tRNA ester bond^81^, providing a plausible alternative mechanism. Both hypotheses center on ester bond cleavage, however, employing different mechanisms. Future high-resolution structural studies will help determine how H44 is oriented relative to the ester bond within ribosome-bound substrates, and whether additional residues contribute to transition state stabilization.

Structural studies will further help elucidate the positioning of Jlp2 and peptidyl-tRNA in the ribosome. At ∼24 kDa - similar to tRNA - Jlp2 is small enough to fit within 80S structures. However, the limited space in or near the PTC would require either local rRNA rearrangements, or peptidyl-tRNA retraction, as recently proposed for Pth^62^ to allow Jlp2’s catalytic cavity access to the peptidyl-tRNA ester bond, potentially explaining why Jlp2 does not act on unsplit/translating ribosomes.

H44 is conserved across Jlp2 orthologs, including the mammalian CCDC25 (Supplementary Fig. S3a). While Jlp2 is cytoplasmic^73^, mammalian CCDC25 also has plasma-membrane-localized pools proposed to promote cancer metastasis by sensing neutrophil extracellular trap DNA, mediated by the NFACT-R domain^82^. Since the CCDC25 NFACT-R domain harbours a histidine (H41) orthologous to H44, it will be important to determine if CCDC25 displays release factor activity, whether this function is conserved across other Jlp2 orthologs, and whether it is linked to CCDC25’s role in cancer metastasis. Rqc2, the only other yeast protein sharing the NFACT-R domain, does not contain histidine at this position. Whether other proteins harboring an NFACT-R domain with an analogous histidine share release activity and whether histidine containing NFACT-R domains constitute a novel class of release factor domains remains to be determined.

In summary, our work defines Jlp2 as a novel factor of an alternative TQC pathway in which peptide release from aberrant peptidyl-tRNA occurs independently of the RQC complex and of previously characterized release factors. By releasing aberrant nascent peptides from ribosome-bound tRNA, Jlp2 restricts pathological CAT tail extension, and independently protects cells against the detrimental consequences of translation elongation stress, revealing an unanticipated layer of complexity in the eukaryotic quality control network.

## METHODS

### Materials

Materials, strains, oligonucleotides and recombinant DNA used in this study are listed in Supplementary Table S1.

### Experimental Model and Subject Details

Strains used in this study were *S. cerevisiae* BY4741 (*MATa his3Δ1 leu2Δ0 met15Δ0 ura3Δ0*)^83^. The genotypes of the strains and their mutant derivatives are listed in Supplementary Table S1. Cells were grown in yeast extract peptone dextrose (YPD) or synthetic dropout medium at 30°C with shaking at 220 RPM. The dropout media contained either 2% glucose (SD) or in the case of PrANS reporter containing cells, 2% raffinose (SRaf). For galactose induction, cells were grown in SRaf medium and 20% galactose was added to a final concentration of 1.8% for 2 hours before harvesting.

### Method Details

#### Yeast strains

Chromosomal manipulations of yeast strains including deletions and tagging were performed by homologous recombination using PCR amplified DNA cassettes^84–86^. PCR was performed using Phusion High-Fidelity DNA Polymerase. Transformation of DNA cassettes and plasmids into yeast was performed using the LiAc/ssDNA/PEG method^87^. Yeast strains used are listed in Supplementary Table S1.

#### Plasmid construction

PCR amplifications of the inserts were performed using Phusion High-Fidelity DNA Polymerase. For cloning of pGFP-TEV-Rz, GFP-TEV was amplified from pGTRR^17^ with primers containing XbaI sites and the hammerhead ribozyme sequence and cloned into the pGTRR^17^ backbone between the XbaI restriction sites. For cloning of Jlp2 overexpressing plasmids, PCR amplification of the *JLP2* coding region excluding the stop codon was performed using the genomic DNA of yeast as the template. Primers used for the amplification contained overhangs for the XbaI restriction site on the forward and reverse primers and additionally the C-terminal FLAG or HA tag followed by a stop codon in the reverse primer. Point mutants were generated by overlap PCR^88^ using primers containing the desired mutations. The *JLP2-HA* insert was cloned into the pGTRR^17^ backbone between the XbaI sites. The *JLP2-FLAG* insert was cloned into the backbone of both pGTRR^17^ and pKK148^37^ and the *JLP2-FLAG* point mutants were cloned into the backbone of pKK148^37^ between the XbaI sites. Primers used are listed in Supplementary Table S1.

DNA cloning was performed by restriction digestion using XbaI and ligation using T4 DNA ligase as per manufacturer’s instructions. Ligation reactions were transformed into chemically competent DH5ɑ *E. coli* cells, selected on LB agar plates containing 100 µg/mL ampicillin and verified by Sanger sequencing.

#### Spot assay

Yeast strains were grown overnight either in YPD or SD media. The following day, cells were diluted to an OD_600_ of 1.0 in media. Four 10-fold dilutions were performed to obtain a total of five different dilutions in media. 10 µL spots from each dilution were spotted next to each other on plates containing the indicated final concentration of cycloheximide (CHX) or Anisomycin (ANI). The plates were then incubated at 30°C.

#### Preparation of lysates and Western blotting

For analysis of CAT tailing and protein levels, whole cell extracts were prepared from log phase cells by chemical lysis method^89^. Briefly, cell pellets were resuspended in 200 µL chemical lysis solution (0.1 M NaOH, 0.05 M EDTA, 2% SDS and 2% (v/v) 2-mercaptoethanol) and boiled for 10 min at 95°C. The lysate was then vortexed at maximum speed followed by addition of 5 µL of 4 M acetic acid and (4x) Laemmli buffer to a final concentration of 1x. The lysates were vortexed and boiled for 10 min at 95°C. The lysates were cleared by centrifugation at maximum speed (18,620 xg) for 5 min at room temperature and the supernatant was transferred to a fresh tube. When CAT tailing and aggregates were analyzed, clearing was performed at 100 xg for 5 min. For TEV treatment, the bead-beating method of lysis was used. Briefly, cell pellets were resuspended in 300 µL of WB lysis buffer (20 mM HEPES-KOH pH 7, 100 mM potassium acetate and 5 mM magnesium acetate). ∼300 µL of zirconia beads were added and cells were intermittently vortexed vigorously for 30 s and placed on ice for 30 s for a total of 10 cycles. The lysates were cleared by centrifugation at 100 xg for 5 min at 4°C and 100 µL of supernatant was transferred to a fresh tube. 1 µL of TEV protease was added per 40 µL of lysate and incubated at 18°C for 2 h with shaking prior to the addition of Laemmli buffer and boiling at 95°C. Lysates were run on 10% SDS-PAGE gels.

For analysis of peptidyl-tRNA, the bead-beating method was used with modifications. Briefly, cell pellets were resuspended in 300 µL of WB lysis buffer along with the addition of 10% SDS and lysed as described above but at room temperature. The lysates were cleared by centrifugation at 18,620 xg for 5 min and 100 µL of supernatants were transferred to a fresh tube. 1 µL of 4M acetic acid was added prior to the addition of (4x) Laemmli buffer to a final concentration of 1x and boiled at 65°C for 5 min. Alternatively, cell pellets were directly resuspended in 300 µL of mixture containing WB lysis buffer, 10% SDS and 1x final conc. Laemmli buffer and boiled using a thermoshaker at 95°C for 5 min with 1500 RPM shaking in the first and last minute of the boiling. The lysates were cleared by centrifugation at 18,620 xg for 5 min and 100 µL of supernatants were transferred to a fresh tube.

For electrophoretic separation, either standard SDS-PAGE or NuPAGE-based (for peptidyl-tRNA analysis) separation was performed. After electrophoresis, proteins were then transferred onto a nitrocellulose membrane and probed with the corresponding antibodies followed by visualization using ECL. The antibodies used are listed in Supplementary Table S1.

#### Northern blot

Northern blotting was performed as described previously^90^. Briefly, 25 mL log phase cultures at OD_600_ were harvested and cells were snap frozen in 750 µL TRIzol Reagent. For RNA extraction, ∼150 µL zirconia/glass beads were added to each tube, vortexed for 30 s and kept on ice for 30 s for a total of 10 cycles. RNA extraction was performed as per manufacturer’s instructions but with an additional acid phenol/chloroform extraction of the aqueous phase. Equal volume of RNA and NorthernMax™-Gly Sample Loading Dye and 0.15 µg/µL additional ethidium bromide were added to the samples and incubated for 30 min at 50°C. 25 µg total RNA from each sample was loaded on a 1.4% w/v agarose gel in MOPS Buffer (200 mM MOPS pH 7, 80 mM sodium acetate, 10 mM EDTA). RNA was transferred using the capillary-based transfer method with 10x SSC (1.5 M sodium chloride, 0.15 M trisodium citrate dihydrate pH 7) onto a nylon membrane. RNA was immobilized on the membrane by crosslinking twice at 1,200 µJoules × 100 doses with the UV Stratalinker™ 1800. The membrane was blocked with hybridization buffer (0.5 M sodium phosphate, 1 mM EDTA and 7 % w/v SDS) and 2 mg salmon sperm DNA for 1 h at 50°C. Blots were probed in hybridization buffer overnight with oligonucleotides targeting *GFP* mRNA, phosphorylated with γ-_32_P-ATP. The membrane was washed three times for up to 45 min with NB washing solution (1 M sodium phosphate, 1 mM EDTA, 1% w/v SDS), and a phosphor screen was exposed to the membrane and visualized with the Typhoon FLA9500 system. The membranes were stripped twice using NB stripping solution (0.5% w/v SDS) for 1 hour at 65°C, blocked and re-probed for *PGK1* mRNA as described above. Probe sequences used are listed in Table S1.

#### Sucrose density gradient fractionation

Cell pellets from 120 mL cultures at 0.8 OD_600_ were resuspended in 400 µL of ice-cold Lysis Buffer S (20 mM HEPES-KOH pH 7.4, 100 mM potassium acetate, 4 mM magnesium acetate, 100 µg/mL cycloheximide and cOmplete EDTA-free protease inhibitor cocktail added just prior to use). All further steps were performed on ice or at 4°C. Approximately 300 µL of zirconia/glass beads were added to the suspensions. The tubes were vortexed at maximum speed for 30 s and placed on ice for 30 s for a total of 10 cycles. The lysates were then transferred to a fresh tube and centrifuged at 18,620 xg for 10 min and the cleared supernatant was transferred to a fresh tube. The supernatant was then layered on top of a 10-45% linear sucrose density gradient in Lysis Buffer S. Ultracentrifugation was performed at 38,000 RPM for 2.5 h using an SW40-Ti rotor. The gradients were then fractionated into ∼0.2 or 0.5 mL fractions using a BIOCOMP Gradient Station. 100 µL of each fraction were then mixed with 1 µL of 4 M acetic acid, prior to the addition of (4x) Laemmli buffer to a final concentration of 1x. The samples were then heated for 10 min at 65°C, before being subjected to western blotting.

#### Co-immunoprecipitation of Jlp2

*jlp2*Δ cells expressing Jlp2-FLAG or Jlp2-HA (negative control) were grown to log phase in 100 mL SD-Ura medium. Cells were harvested and resuspended in 500 µL of co-IP lysis buffer (20 mM HEPES-KOH pH 7, 100 mM potassium acetate, 5 mM magnesium acetate, 0.075% IGEPAL and 1x protease inhibitor cocktail). ∼400 µL of zirconia beads were added to the cell suspension. Cells were intermittently vortexed at 15,620 xg for 30 s and placed on ice for 30 s for a total of 10 cycles. Another 500 µL of co-IP lysis buffer were added and the lysate was transferred to a fresh tube and cleared by centrifugation at 15,620 xg for 10 min at 4°C. The cleared supernatant was transferred to a fresh tube containing 25 µL anti-FLAG magnetic beads pre-washed with co-IP lysis buffer and incubated at 4°C for 45 min on a rotating wheel. The beads were then washed 3 times with co-IP lysis buffer on the rotating wheel at 4°C for 5 min each. The supernatant was removed, and the beads were incubated with 40 µL of 1x NuPAGE loading dye diluted in MilliQ water and heated at 60°C for 10 min. The supernatant was then transferred to a fresh tube and used for MS analysis.

#### Purification of Jlp2

The appropriate knockout strains (corresponding to the knockouts from which PrANS peptidyl-tRNA was enriched for *in vitro* reconstitution) containing *pJLP2-FLAG* or the substitution mutants were grown overnight in SD-His media and diluted the next day to an OD_600_ of 0.2 in 500 mL of SD-His media. Cells were harvested at an OD_600_ of 0.8 by centrifugation of cultures at 4000 xg for 3 min. The supernatant was discarded, and the cell pellets were washed with MilliQ water and centrifuged again at 5340 xg for 1.5 min. The supernatant was discarded, the cell pellet was flash frozen in liquid nitrogen and stored at -20°C. Cell pellets were thawed on ice after adding 500 µL Lysis buffer P (20 mM HEPES-KOH pH 7.4, 100 mM potassium acetate, 10 mM magnesium acetate and cOmplete EDTA-free protease inhibitor cocktail added just prior to use). All further steps were performed on ice or at 4°C. Approximately 500 µL of zirconia beads were added to the suspensions and the tubes were vortexed at maximum speed for 30 s and placed on ice for 30 s for a total of 10 cycles. The lysates were transferred to fresh 2mL tubes and an additional 1 mL of Lysis Buffer P was added to the lysates and centrifuged at 18,620 xg for 10 min. The cleared supernatant was transferred to a fresh tube containing 100 µL of Anti-FLAG® M2 Magnetic Beads slurry pre-washed 3 times in Lysis Buffer P for 5 min. The tubes were then incubated for 2 h on a rotating wheel, then placed on a magnetic rack and the supernatant was removed and discarded. The magnetic beads were washed thrice with 1.5 mL Lysis Buffer P for 5 min on the rotating wheel. Placing the tubes on the magnetic rack, the supernatants were discarded and 1.5 mL Lysis Buffer H (20 mM HEPES-KOH pH 7.4, 800 mM potassium acetate, 10 mM magnesium acetate and cOmplete EDTA-free protease inhibitor cocktail added just prior to use) was added to the tubes and incubated for 10 min on the rotating wheel. Two more washes with Lysis Buffer H were performed, followed by three washes with Lysis Buffer P. The supernatant was discarded, and the beads were resuspended in 400 µL of Lysis Buffer P containing 450 µg/mL FLAG peptide for elution of Jlp2-FLAG, Jlp2(H44A)-FLAG or Jlp2(H44F)-FLAG. Elution was performed for 1 hour on the rotating wheel. The eluate was then transferred to a fresh tube. Buffer exchange to Jlp2 Storage Buffer (20 mM HEPES-KOH pH 7.4, 100 mM potassium acetate, 10 mM magnesium acetate and 10% glycerol) and removal of excess FLAG peptide was performed using Vivaspin 2 (MWCO 5 kDa) or Amicon® Ultra (10 kDa MWCO) centrifugation filters as per manufacturer’s instructions until fresh buffer was >95%. The purified proteins were stored at -20°C.

#### Sucrose density gradient-based enrichment of peptidyl-tRNA:ribosome complexes for *in vitro* reconstitution

The appropriate yeast strains containing *pAV184* were grown overnight in SRaf-Ura media. The next day, cultures were diluted to an OD_600_ of 0.2 in 500 mL of SRaf-Ura media and grown to an OD_600_ of 0.6. Reporter protein expression was induced with 1.8% (final conc.) galactose for 2 h and the cells were harvested by centrifugation at 4000 xg for 3 min. The supernatant was discarded, and the cell pellets were washed with MilliQ water and centrifuged again at 5340 xg for 1.5 min. The supernatant was discarded, and the cell pellet was flash frozen in liquid nitrogen and stored at -20°C. Peptidyl-tRNA ribosome complexes were enriched by sucrose density gradient fractionation (as described above with the following modifications) under either “Physiological Salt” buffer (PS) conditions (20 mM HEPES-KOH pH 7, 100 mM potassium acetate, 10 mM magnesium acetate and cOmplete EDTA-free protease inhibitor cocktail added just prior to use) or “High Salt” buffer (HS) conditions (20 mM HEPES-KOH pH 7, 750 mM Potassium acetate, 10 mM magnesium acetate and cOmplete EDTA-free protease inhibitor cocktail added just prior to use) for both Lysis and Gradient Buffers. The salt concentrations used for the “physiological salt” condition are within the ranges commonly employed in sucrose density gradient fractionation^9,15,24,31^ and *in vitro* translation^91,92^ experiments. 60S and 80S fractions were collected separately, and buffer exchange to Ribosome Storage Buffer (20 mM HEPES-KOH pH 7, 100 mM potassium acetate, 10 mM magnesium acetate and 10% glycerol) was performed as per manufacturer’s instructions. Preparations were stored at -20°C.

Enriched peptidyl-tRNA:ribosome complexes used for *in vitro* reconstitution were quantified via with the NanoDrop 2000 system in RNA mode and an amount equivalent to 6000 ng of RNA was mixed with 6 µM (final concentration) of purified Jlp2-FLAG in a 60 µL reaction and incubated at 30°C for 20 min. The reaction was stopped by the addition of (4x) Laemmli buffer to a final concentration of 1x and heated to 65°C for 5 min. The resulting samples were used for Western blotting.

#### IgG magnetic bead-based enrichment of peptidyl-tRNA:ribosome complexes for on-bead *in vitro* reconstitution

The appropriate yeast strains containing *pAV184* were grown and reporter protein expression was induced, followed by harvesting as described above. Cell pellets were thawed on ice after adding 500 µL Lysis Buffer PS. All further steps were performed on ice or at 4°C. Approximately 500 µL of zirconia beads were added to the suspensions and the tubes were vortexed at maximum speed for 30 s and placed on ice for 30 s for a total of 10 cycles. The lysates were transferred to fresh 2mL tubes and an additional 1 mL of Lysis buffer PS was added to the lysates and centrifuged at 18,620 xg for 10 min. The cleared supernatant was transferred to a fresh tube containing 180 µL of Rabbit (DA1E) mAb IgG XP® Magnetic Bead Conjugate slurry that were pre-washed 3 times in Lysis Buffer P for 5 min. The tubes were then incubated for 2 h on a rotating wheel, then placed on a magnetic rack and the supernatant was removed and discarded. The magnetic beads were washed thrice with 1.5 mL Lysis Buffer P for 5 min on the rotating wheel. With the final wash step, the suspension was equally split into two sets (1 and 2 respectively). Placing the tubes on the magnetic rack, the supernatants were discarded and 1.5 mL of either Lysis Buffer P (Set 1 – ‘physiological salt’) or Lysis Buffer H (Set 2 – ‘high salt’) was added to the two sets of tubes and incubated for 10 min on the rotating wheel.

Two more washes with either Lysis Buffer P or H were performed for Set 1 and 2, respectively, followed by three washes with Lysis Buffer P for both sets. The supernatant was discarded, and the beads were resuspended in 120 µL of Lysis Buffer P. The suspensions were then separated into 40 µL aliquots and mixed with purified Jlp2-FLAG (5 µM final conc.) in a 60 µL reaction and incubated at 30°C for 20 min. The reaction was stopped by the addition of (4x) Laemmli buffer to a final concentration of 1x and heating to 65°C for 5 min at the indicated time points. This step also allowed for the release of protein A (from the nascent peptides) from the IgG beads. The supernatants were transferred to a fresh tube and used for Western blotting using NuPAGE gels. For analysis of peptidyl-tRNA 60S and 80S complexes separately, the corresponding complexes were first enriched using sucrose density gradient fractionation with minor modifications. Specifically, for enrichment of 80S complexes, sucrose density gradient fractionation was performed using Buffer PS as described above. For enrichment of 60S, magnesium acetate was omitted from Buffer PS to dissociate all 80S ribosomes to enrich for 60S complexes and avoid re-association. The corresponding fractions were then used for immunoprecipitation using IgG magnetic beads followed by salt washes and *in vitro* reconstitution as described above.

### LC-MS/MS analysis

#### LC-MS/MS analysis was performed as previously described^90^

For the identification of Jlp2 interactors, LC-MS/MS analysis was performed on four replicates, each, of Jlp2-FLAG and negative control (Jlp2-HA, which does not bind to anti-FLAG beads but expected to replicate overall cellular proteostasis as in Jlp2-FLAG) co-immunoprecipitation samples.

#### Enzymatic protein digestion

All samples were processed using the SP3 approach^93^. Proteins were reduced with 5 mM DTT, alkylated with 15 mM iodoacetamide in the dark and quenched with 5 mM DTT. Enzymatic protein digestion was performed using trypsin at 37°C overnight. The resultant peptide solution was purified by solid phase extraction in C18 StageTips^94^.

#### Liquid chromatography tandem mass spectrometry

Peptides were separated via an in-house packed 45-cm analytical column (inner diameter: 75 μm; ReproSil-Pur 120 C18-AQ 1.9-μm silica particles, Dr. Maisch GmbH) on a Vanquish Neo UHPLC system (Thermo Fisher Scientific). Online reverse-phase chromatography was performed through a 70-min non-linear gradient of 1.6-32% acetonitrile with 0.1% formic acid at a nanoflow rate of 300 nl/min. The eluted peptides were sprayed directly by electrospray ionization into an Orbitrap Astral mass spectrometer (Thermo Fisher Scientific). Mass spectrometry was conducted in data-dependent acquisition mode using a top50 method with one full scan in the Orbitrap analyzer (scan range: 325 to 1,300 m/z; resolution: 120,000, target value: 3 × 106, maximum injection time: 25 ms) followed by 50 fragment scans in the Astral analyzer via higher energy collision dissociation (HCD; normalized collision energy: 26%, scan range: 150 to 2,000 m/z, target value: 1 × 104, maximum injection time: 10 ms, isolation window: 1.4 m/z). Precursor ions of unassigned, +1 or higher than +6 charge state were rejected. Additionally, precursor ions already isolated for fragmentation were dynamically excluded for 15 s.

#### Mass spectrometry data analysis

Mass spectrometry raw data were processed by MaxQuant software (version 2.6.2.0)^95^ using its built-in Andromeda search engine^96^. MS/MS spectra were searched against a target-decoy database containing the forward and reverse protein sequences of UniProt *S. cerevisiae* reference proteome (release 2024_03; 6,091 entries), the corresponding transgenic proteins and a default list of common contaminants. Carbamidomethylation of cysteine was considered as fixed modification. Methionine oxidation and protein N-terminal acetylation were assigned as variable modifications. A maximum of 2 missed cleavages were allowed. The “match between runs” function was activated. The “second peptides” option was switched on. Minimum peptide length was set to 7 amino acids. False discovery rate (FDR) was set to 1% at both peptide and protein levels.

Protein quantification was performed using the MaxLFQ algorithm^97^ skipping its default normalization option. Minimum LFQ ratio count was set to one. Both the unique and razor peptides were used for quantification. Differential expression analysis was performed in R statistical environment. Reverse hits, potential contaminants and “only identified by site” protein groups were first filtered out. Data normalization was performed on the log-transformed data using median-centering (Jlp2-FLAG vs HA experiment) or based on Protein A intensities followed by median-centering within each group (high salt vs physiological salt experiment). Proteins were further filtered to retain only those detected in at least 3 out of the 4 replicates in either group of each comparison. Following imputation of the missing LFQ intensity values, a linear model was fitted using the limma package in R^98^ to assess the difference between the two groups for each protein, with adjustment for multiple testing using the Benjamini-Hochberg approach^99^. The log2 fold change and the significance of the difference were displayed on a volcano plot. Only proteins with a minimum log2 fold change of 1 and an FDR-adjusted p value (q value) lower than 0.05 were considered differentially regulated.

### Sequence and Structural analysis

Multiple sequence alignment was performed using T-Coffee^63^ Version_11.00, Structural alignment mode (M-Coffee) accessed on 09 Nov 2025 and the results were visualized using JalviewJS^100^ and saved as .aln file. . Conservation scores derived from the multiple sequence alignment were mapped onto the protein structure using ChimeraX^101^. The AlphaFold predicted structure of Jlp2 (AF-P40206-F1-v6) was obtained from AlphaFold protein structure database^52^ and the X-ray crystal structure of human CCDC25 (PDB ID: 7EOE) was obtained from PDB^102,103^. Structures were visualized and manipulated using UCSF ChimeraX^101^. Cavity was defined using pyKVFinder^104^ accessed through the “find cavities” function in ChimeraX^101^. Folding stability analysis for amino acid substitutions were performed using PremPS^105^ accessed on 11 May 2026 and DynaMut2^106^ accessed on 7 May 2026. Co-folding predictions were made using the AlphaFold3^64^ reproduction, OpenFold3^65,66^ (v0.4.1) accessed on 12 Jul 2026 on Tamarind Bio^107^ using the primary sequence of Jlp2 (Uniprot ID P40206) and the CCA-3xAla SMILES file uploaded to ModelArchive^108^ under the model ID: ma-lgusi.

### Signal Quantification and processing

Quantifications for analysis of Western blots were performed using ImageJ^109^ by selecting the bands using the rectangle tool followed by the generation of intensity plots. The area under the curve of the intensity plot for each band was then measured. For figures in panels showing CAT tail-dependent aggregates and peptidyl-tRNA from GFP-Rz reporter, brightness and contrast were adjusted linearly and uniformly across the entire image shown in the box using ImageJ^109^ for better visualization.

### Statistical Analysis

Student’s t-test was performed using Microsoft Excel and the type of Student’s t-test used for analysis is indicated in the corresponding figure captions. The p-values are shown on the graphs above the corresponding bars and connected by lines showing the samples compared. A p-value of 0.05 or below was generally considered significant. For proteomics data, statistical analysis was performed as described above using R statistical environment.

## Resource availability

### Lead contact

Further information and requests for resources and reagents should be directed to and will be fulfilled by the Lead Contact, Marie-Luise Winz.

### Materials availability

All unique/stable reagents generated in this study are available from the Lead Contact.

## Data Availability

The mass spectrometry proteomics data have been deposited to the ProteomeXchange Consortium via the PRIDE^110^ partner repository with the dataset identifier PXD067959.

The OpenFold3-predicted model structures reported in this study have been deposited in ModelArchive^108^ under the model ID: ma-lgusi

## Supplementary Information

Supplementary Figures S1-S5

Supplementary Table S1. LC-MS/MS analysis for Jlp2 co-immunoprecipitation, related to Supplementary Figure S2

Supplementary Table S2. Predicted ΔΔG values for Jlp2 H44, H52 and N27 mutants and their orthologous mutations in hsCCDC25.

Supplementary Table S3. Reagents, yeast strains, oligonucleotides and recombinant DNA used in this study

## Acknowledgements

M.L.W. acknowledges funding by Deutsche Forschungsgemeinschaft (DFG, German Research Foundation) [project number 439669440 TRR319 RMaP TP B05 and TP B07 to M.L.W., Project number 255344185 SPP1784, Startup Funding to M.L.W.] and by the Research Initiative Rhineland-Palatinate [Startup Funding by the ReALity Initiative to M.L.W.]. Funding from the DFG also supported the Orbitrap Astral system (DFG Project number 524805621; Proteomics Core Facility at IMB-Mainz). We gratefully acknowledge Brian Luke (Johannes Gutenberg University Mainz) and Helle Ulrich (Johannes Gutenberg University Mainz) for plasmids that were used for the generation of deletion strains. We are grateful to Kamena Kostova (Stowers Institute for Medical Research), Jonathan Weissman (University of California, San Francisco) and Ambro van Hoof (University of Texas Health Science Center at Houston) for sharing important reporter constructs used in this study. We acknowledge and thank the Proteomics Core Facility at Institute of Molecular Biology (IMB) Mainz and its members-Jiaxuan Chen, Amitkumar Fulzele, Mario Dejung and Jasmin Cartano, for proteomics analysis including sample processing, mass spectrometry measurement and data analysis. We thank Kristina Friedland (Johannes Gutenberg-University Mainz) and Jean-Yves Roignant (Université de Lausanne) for sharing instruments and reagents used in this work. We also acknowledge Simone Zimmerman, Sophie Mulartschyk, Fabienne Helmes-Stumpf, and Laura Sophie Hartmann (all Johannes Gutenberg-University Mainz) and Zeynep Buse Orhan (Izmir Institute of Technology) for assistance in biochemical experimentation, as well as Christian Kersten (Johannes Gutenberg-University Mainz) for helpful discussions on structure predictions and modelling.

## Contributions

M.L.W. and K.V.I. conceived the study; K.V.I. and M.L.W. designed experiments; K.V.I., A.A.K. and G.F.S. performed and analysed Western blot experiments; A.A.K. performed spot assay experiments; K.V.I. and A.A.K. constructed strains; K.V.I. and C.A.W. performed and analysed sucrose density gradient fractionation experiments; K.V.I. and M.M. performed Cloning experiments; M.M. performed the Northern Blot experiment; L.S.T. performed co-immunoprecipitation for LC-MS/MS experiments; K.V.I. and M.L.W. analysed proteomics data; K.V.I. analysed evolutionary conservation; K.V.I. and M.L.W. performed and analysed structure predictions; K.V.I. performed and analysed *in vitro* reconstitution experiments; K.V.I. drafted the first version of the manuscript, with input from M.L.W. All authors commented on the manuscript and discussed the data.

