## Supplementary Information for "Jlp2 is a non-canonical release factor that protects cells during ribosome stalling"

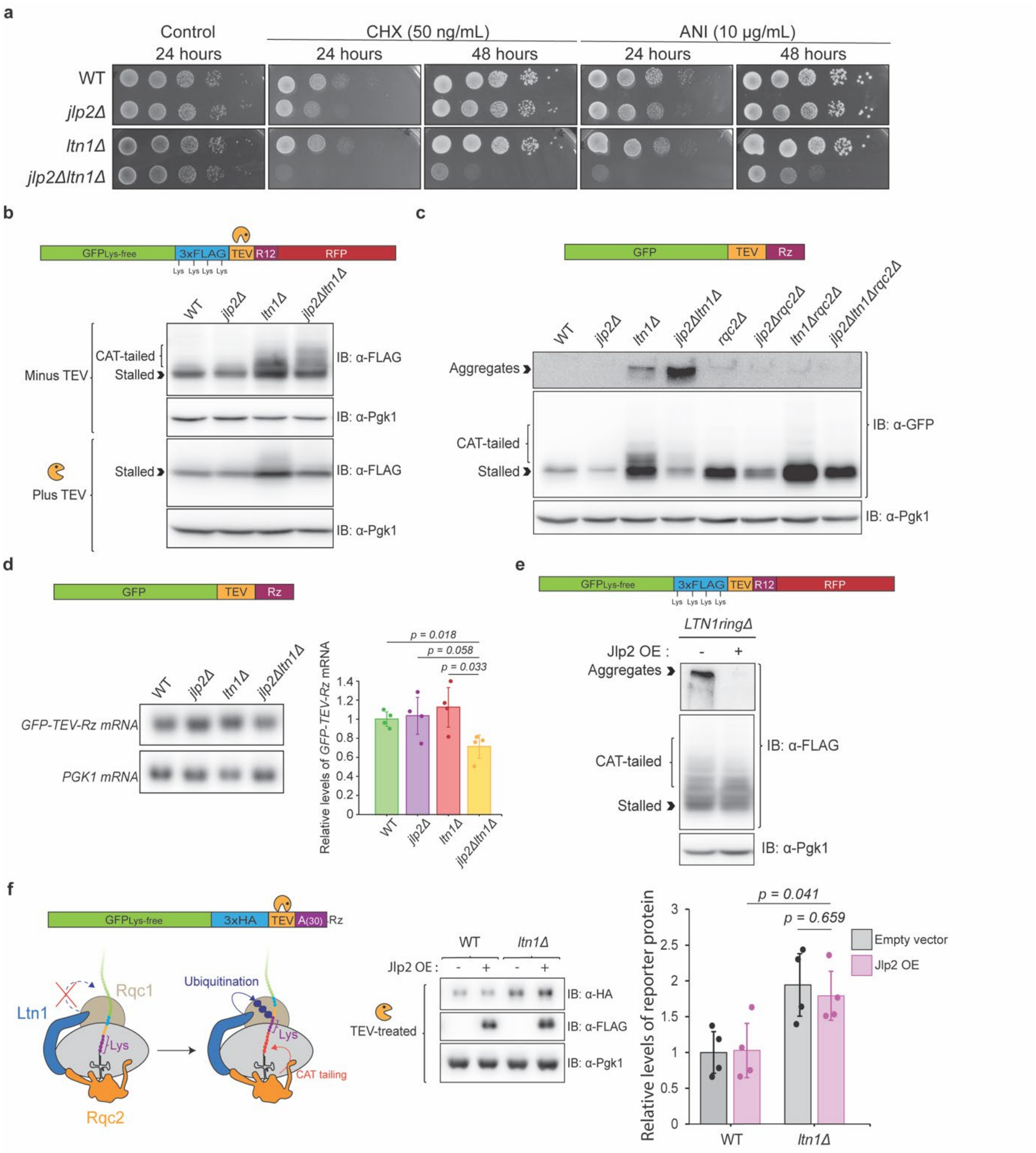


**Supplementary Figure S1. Jlp2 is involved in the quality control of substrates arising from ribosome collisions**

a. *JLP2* genetically interacts with members of the RQC complex. Growth of WT, *jlp2Δ, ltn1Δ* and *jlp2Δltn1Δ* cells spotted in serial dilutions on YPD control and plates containing 50 ng/mL CHX and 10 µg/mL ANI and imaged at the indicated timepoints. Growth defects result in smaller colonies relative to WT.

b. Western blot analysis of TEV protease treated lysates from WT, *jlp2Δ*, *ltn1Δ*, and *jlp2Δltn1Δ* cells expressing GFP$_{Lys\text{-}free}$-3xFLAG-TEV-R12-RFP reporter. Reporter used and TEV cleavage site is indicated at the top of the panel. TEV cleavage collapses protein products of different sizes (different stall positions, +/- CAT tails of heterogeneous length) into a single band, showing that molecular weight shift is a consequence of C-terminal extensions.

c. Deletion of *JLP2* and *LTN1* results in an increase in CAT tail-dependent aggregates from a non-stop truncated reporter. Western blot analysis of GFP-TEV-Rz reporter in WT, *jlp2Δ*, *ltn1Δ*, *jlp2Δltn1Δ*, *rqc2Δ*, *jlp2Δrqc2Δ*, *ltn1Δrqc2Δ* and *jlp2Δltn1Δrqc2Δ*.
d. Northern blot analysis and quantification of *GFP-TEV-Rz* reporter mRNA in WT, *jlp2Δ*, *ltn1Δ* and *jlp2Δltn1Δ*. Reporter mRNA levels were normalized to *PGK1* mRNA. Student's t-test (two-tailed, heteroscedastic) was used to calculate the p-value (N=3 biological replicates, n=1 or 2 technical replicates).
e. Western blot analysis of $GFP_{Lys\text{-}free}$-3xFLAG-TEV-R12-RFP reporter in *LTN1-ringΔ* cells containing either an empty vector or a Jlp2-HA overexpression vector.
f. Jlp2 overexpression does not completely inhibit CAT tailing. Schematic detailing the mechanism of reporter peptide ubiquitination (left). The reporter encodes lysines only at the very C-terminus just upstream of the ribozyme cleavage site (stalling site), and requires CAT tailing for exposure of those lysines for Ltn1-mediated ubiquitination and consecutive proteasomal degradation. A complete inhibition of CAT-tailing in WT upon overexpression of Jlp2 would stabilize the reporter peptide to levels seen in *ltn1Δ* cells. Western blot analysis of $GFP_{Lys\text{-}free}$-3xHA-TEV-A(30)-Rz reporter (TEV cleavage site indicated in the reporter scheme) in WT and *ltn1Δ* cells containing either an empty vector or a Jlp2-FLAG overexpression vector and quantification. Lysates were TEV treated to collapse heterogeneously sized peptide products (+/- CAT-tail, heterogeneous stalling patterns) into a single band for more robust quantification. Student's t-test (two-tailed, heteroscedastic) was used to calculate the p-value (N=3 biological replicates, n=1 or 2 technical replicates).

Schematic representations of the reporter constructs are shown at the top of the corresponding panels.

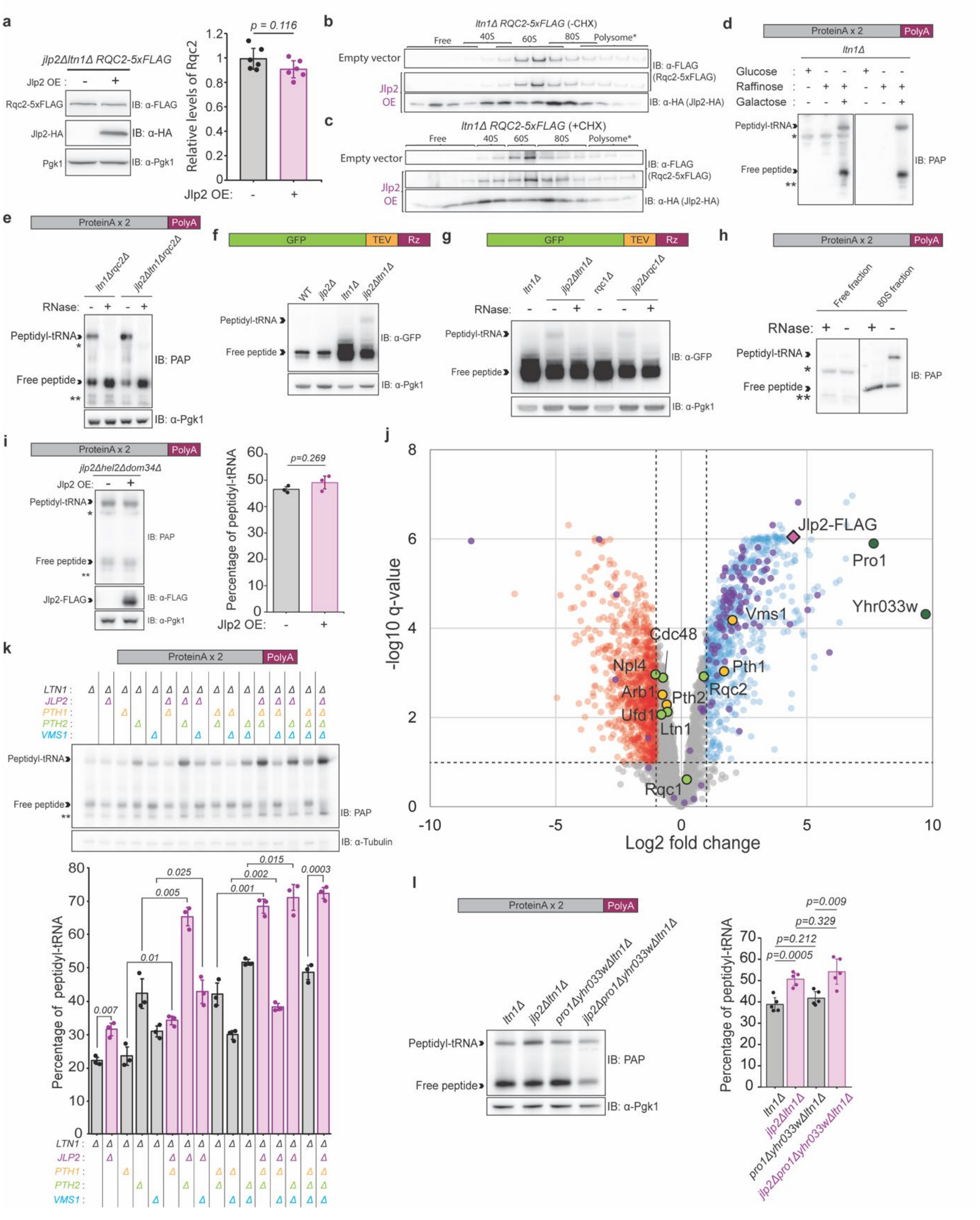

**Supplementary Figure S2. Jlp2 is required for the release of peptidyl-tRNA of ribosome stalling substrates in the absence of canonical RQC**

a. Jlp2 does not regulate Rqc2 expression levels. Western blot analysis of genomically tagged endogenous *RQC2-5xFLAG* in *jlp2Δltn1Δ* cells without and with overexpressing Jlp2-HA and quantification. Reporter protein levels were normalized to Pgk1. Student's t-test (two-tailed, heteroscedastic) was used to calculate the p-value (N=3 biological replicates, n=2 technical replicates).

b. Jlp2 does not compete with Rqc2 for binding to the 60S. Sucrose density gradient fractionation of *ltn1Δ* cells expressing endogenously 5xFLAG-tagged Rqc2 containing either an empty vector or a Jlp2-HA overexpression vector followed by Western blot analysis of the fractions. The experiment was performed without the use of CHX. Asterisk (*) on the Polysome label indicates that only the first four polysome fractions are shown in the figure. Lanes 1 and 2 contain pooled fractions 1+2 and 3+4 while all other following lanes contain fractions 5, 6, etc. Overlapping brackets above some lanes indicate fractions between two peaks containing particles from both.

c. Same as b, except that the cells were treated with 100 µg/mL CHX for 5 min before harvesting.

d. PrANS reporter-specific and non-specific bands observed with the use of the PAP antibody. Western blot showing an inconsistently observed ~40kDa non-specific band (*) and reporter specific bands including peptidyl-tRNA, free peptide and unassigned lower molecular weight PrANS reporter-specific species (**) observed with the use of the PAP antibody. The left and right panels show two independent experiments using the same PrANS reporter- expressing *ltn1Δ* strain grown in either glucose (repression of GAL promoter driving the expression of the PrANS reporter), Raffinose (neither induction nor repression of PrANS reporter expression) or Raffinose and Galactose (strong induction of PrANS reporter expression).

e. Western blot analysis of peptidyl-tRNA species of PrANS reporter in *ltn1Δrqc2Δ* and *jlp2Δltn1Δrqc2Δ* without and with RNase A/T treatment. Asterisk (*) denotes a non-reporter specific (Extended Data Fig. 2d), RNase-resistant (Extended Data Fig. 2h) band observed inconsistently across experiments and double-asterisk (**) denotes unassigned reporter-specific signals (Extended Data Fig. 2d).

f. Western blot analysis of peptidyl-tRNA species of GFP-TEV-Rz reporter in WT, *jlp2Δ, ltn1Δ* and *jlp2Δltn1Δ*.

*g.* Western blot analysis of peptidyl-tRNA and free peptide species of GFP-TEV-Rz reporter in *ltn1Δ, jlp2Δltn1Δ, rqc1Δ* and *jlp2Δrqc1Δ,* without and with RNase A/T treatment in *jlp2Δltn1Δ,* and *jlp2Δrqc1Δ.*

h. RNase A/T treatment confirming the RNase-insensitivity of the nonspecific (Fig. 2d denoted by "*") band observed in the ribosome-free fraction during sucrose density gradient fractionation and Western blot analysis of peptidyl-tRNA species of PrANS reporter in *rqc1Δ* from Figure 2b. Samples were run on the same gel and transferred to the same membrane. Line in the middle indicates that several lanes were spliced out from the original membrane image. Double-asterisk (**) denotes unassigned reporter-specific signals (Extended Data Fig. 2d).

i. Western blot analysis of peptidyl-tRNA and free peptide species of PrANS reporter in *jlp2Δhel2Δdom34Δ* cells containing either an empty vector or Jlp2 overexpression vector and quantification of percentage of peptidyl-tRNA relative to the total peptide. Student's t-test (two-tailed, heteroscedastic) was used to calculate the p-value. N=3 biological replicates. Asterisk (*) denotes a non-reporter specific band (and double-asterisk (**) denotes unassigned reporter-specific signals (Extended Data Fig. 2d).

j. Volcano plot of proteins co-immunoprecipitated with Jlp2-FLAG and identified via label-free quantification by LC-MS/MS. $\log_2$-fold changes and $-\log_{10}$ q-values are plotted on the X and Y axes respectively. Proteins of interest are highlighted: Jlp2-FLAG (magenta), ribosomal proteins (purple), RQC and Cdc48 complex proteins (green), known or predicted release factors (yellow) and Pro1 and Yhr033w (dark green). N=4 biological replicates.

k. Western blot analysis of peptidyl-tRNA species of PrANS reporter in *ltn1Δ* strains additionally lacking *JLP2, PTH1, PTH2* and/or *VMS1* in different combinations and quantification of percentage of peptidyl-tRNA. Student's t-test (two-tailed, heteroscedastic) was used to calculate the p-value. N=3 biological replicates. Double-asterisk (**) denotes unassigned reporter-specific signals (Extended Data Fig. 2d).

l. Western blot analysis of peptidyl-tRNA species of PrANS reporter in *ltn1Δ, jlp2Δltn1Δ, pro1Δyhr033wΔltn1Δ* and *jlp2Δpro1Δyhr033wΔltn1Δ* and quantification of percentage of peptidyl-tRNA. Student's t-test (two-tailed, heteroscedastic) was used to calculate the p-value. N=3 biological replicates, n=1 or 2 technical replicates.

Schematic representations of the reporter constructs are shown at the top of the corresponding panels

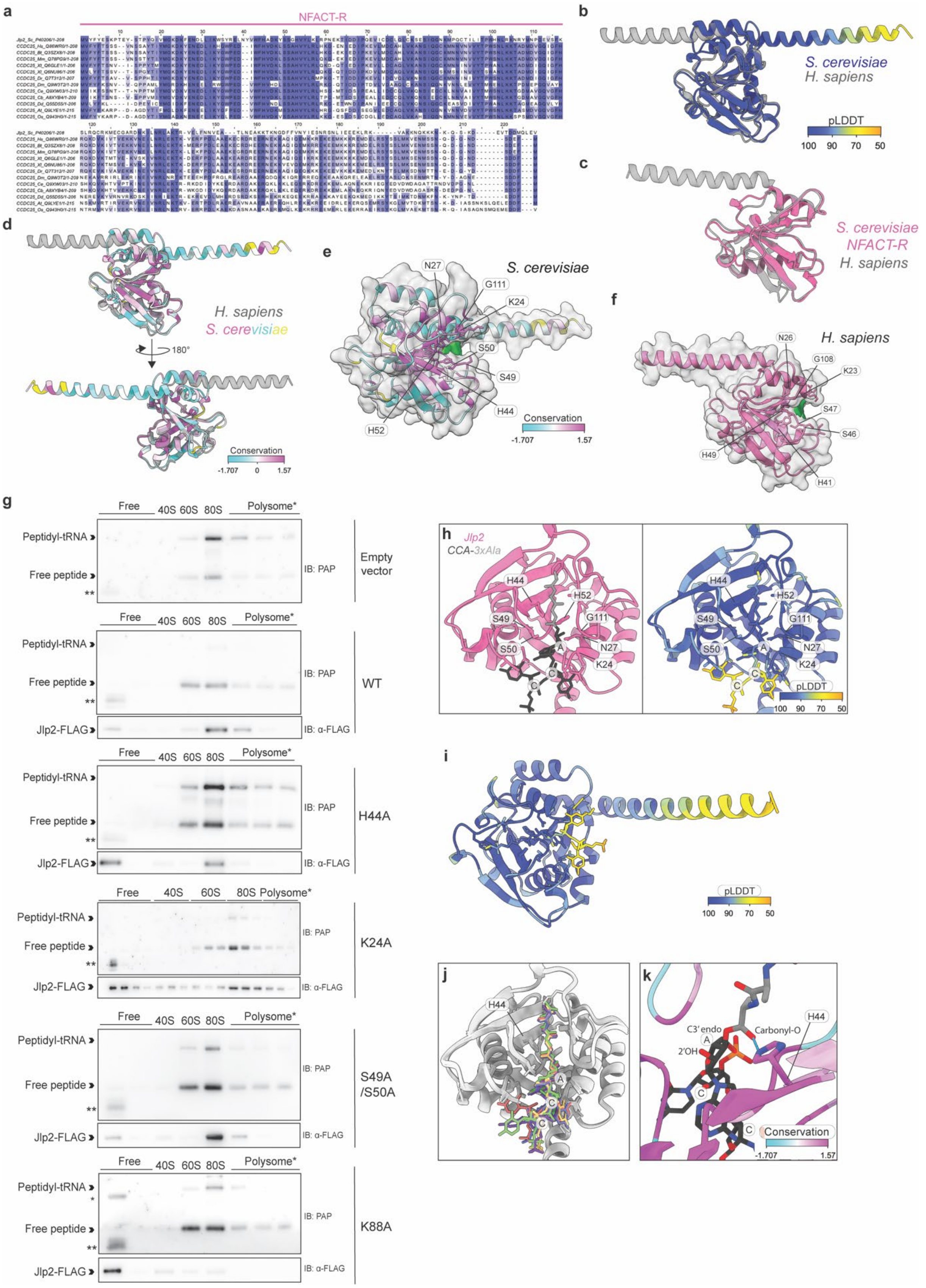


**Supplementary Figure S3. Identification of a conserved residue essential for the release factor activity of Jlp2 using sequence and structure conservation analysis**

a. Evolutionary sequence conservation of Jlp2 across species. Multiple sequence alignment of Jlp2 and its orthologs from the indicated organisms (animals and plants), generated using T-Coffee[63]. Darker shades of purple indicate a higher degree of conservation.
b. Structural conservation between yeast Jlp2 and human CCDC25. The AlphaFold-predicted structure of Jlp2 (AF-P40206-F1-v6)[52], colored according to pLDDT values, superimposed on the X-ray crystal structure of human CCDC25 (PDB: 7EOE, Chain A; grey).
c. Structural conservation of the NFACT-R domain between Jlp2 and human CCDC25. The NFACT-R domain of Jlp2 (residues 1-116) from the AlphaFold-predicted structure (AF-P40206-F1-v6)[52] (pink) superimposed on human CCDC25 (PDB: 7EOE, Chain A; grey).
d. Structural conservation between yeast Jlp2 and human CCDC25. The AlphaFold-predicted structure of Jlp2 (AF-P40206-F1-v6)[52] colored according to evolutionary sequence conservation (scale shown) superimposed on human CCDC25 (PDB: 7EOE, Chain A; grey). Residues shown in yellow correspond to positions for which sequence conservation values could not be determined.
e. A prominent cavity lined by highly conserved residues is present in the NFACT-R domain of Jlp2. The AlphaFold-predicted structure of Jlp2 (AF-P40206-F1-v6)[52] colored according to evolutionary sequence conservation (scale shown), displayed with its predicted molecular surface (transparent grey). The cavity is indicated by green density. Residues shown in yellow correspond to positions for which sequence conservation values could not be determined.
f. An orthologous cavity is present in human CCDC25. The X-ray crystal structure of human CCDC25 (PDB: 7EOE, Chain A; pink) displayed with its molecular surface (transparent grey). The cavity corresponding to that identified in Fig. 3a and Extended Data Fig. 3e is indicated by green density.
g. Peptide release activity and ribosomal association of selected Jlp2 mutants. Sucrose density gradient fractionation followed by Western blot analysis of PrANS reporter peptidyl-tRNA species and the ribosomal distribution of the indicated Jlp2 mutants in *jlp2Δltn1Δrqc2Δ* cells. Double-asterisk (**) denotes unassigned, reporter-specific signals (Extended Data Fig. 2d).
h. Zoomed-in view of OpenFold3-predicted model of Jlp2 associated with CCA-3xAla ligand (Model 1). Left: Jlp2 (pink) with CCA (black) - 3xAlanine (grey), right: Model on left colored according to per-residue pLDDT. Residues at the predicted interface are labelled. Model confidence metrics are IPTM = 0.856047, PTM = 0.885398, average PLDDT = 90.754242, GPDE = 0.671759 and Sample ranking score = 0.895571.

i. Highest-ranking OpenFold3-predicted model of Jlp2 associated with CCA-3xAla ligand (Model 1) colored according to pLDDT scores.
j. All five OpenFold3-predicted models of the Jlp2–CCA-3×Ala complex, superimposed, showing consistent positioning of the ligand within the NFACT-R cavity across independent predictions. Jlp2 is shown in light grey; the CCA-3×Ala ligand is coloured by model as follows. Model 1 (red) IPTM = 0.856047, PTM = 0.885398, average PLDDT = 90.754242, GPDE = 0.671759 and Sample ranking score = 0.895571, 2 (yellow) IPTM = 0.790356, PTM = 0.86498 , average PLDDT = 90.199638, GPDE = 0.808211 and Sample ranking score = 0.843742, 3 (green) IPTM = 0.820688, PTM = 0.872538, average PLDDT = 90.3858, GPDE = 0.712759 and Sample ranking score = 0.869519, 4 (blue) IPTM = 0.82805, PTM = 0.874493, average PLDDT = 90.404381, GPDE = 0.741862 and Sample ranking score = 0.870993, 5 (purple) IPTM = 0.828802, PTM = 0.871843, average PLDDT = 89.9477, GPDE = 0.700878 and Sample ranking score = 0.871064.
k. Zoomed-in view of OpenFold3-predicted model of Jlp2 (colored according to sequence conservation as in Fig. 3a) associated with CCA-3xAla (black-grey) ligand (Model 1). Dotted blue line represents a predicted hydrogen bond between H44's ring $N^{\tau}$H and the carbonyl-oxygen of the ester bond. Nitrogen atoms are colored in blue and Oxygen atoms in red.

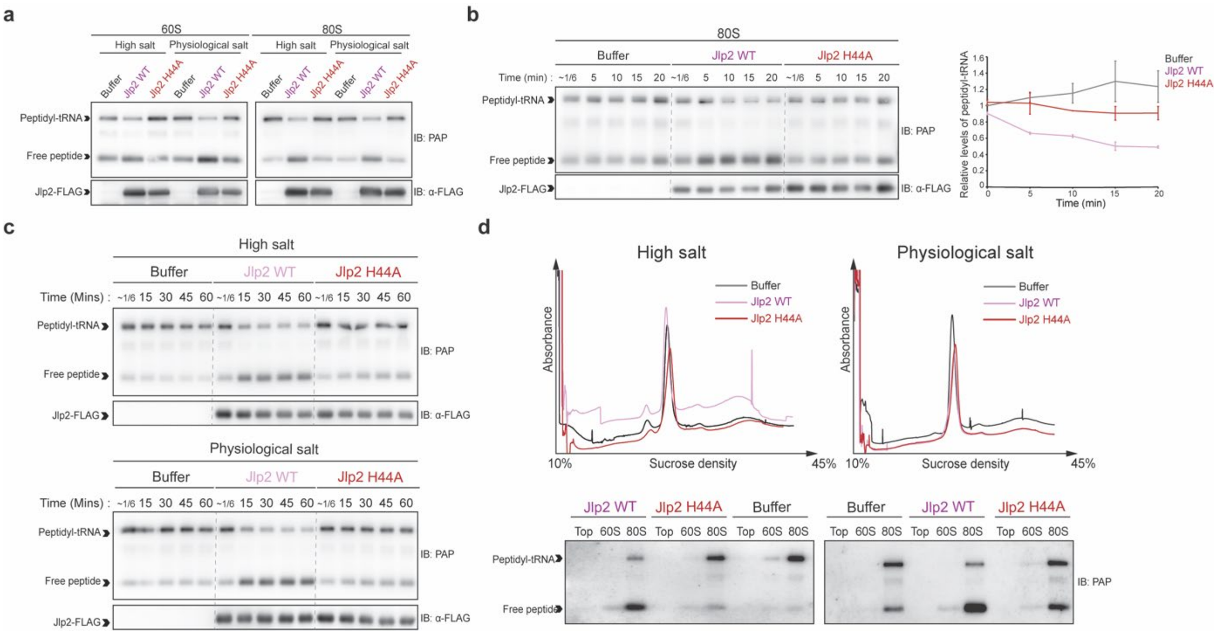


**Supplementary Figure S4. Jlp2 directly catalyzes the release of nascent peptides that escape canonical RQC processing**

a. Wild-type Jlp2 catalyzes peptide release from both 60S and 80S-associated peptidyl-tRNA independently of fractionation conditions. *In vitro* reconstitution using peptidyl-tRNA:60S and peptidyl-tRNA:80S fractions enriched in either high salt or physiological salt conditions by sucrose density gradient fractionation from *jlp2Δltn1Δrqc2Δ* cells with purified WT and H44A mutant Jlp2-FLAG followed by Western blot analysis for peptidyl-tRNA release.

b. Time course analysis of sucrose density gradient-enriched peptidyl-tRNA:80S fractions from *jlp2Δltn1Δrqc2Δ* cells with purified WT and H44A mutant Jlp2-FLAG followed by Western blot analysis for peptidyl-tRNA release and quantification (n=2 independent experiments). For quantification, the percentage of peptidyl-tRNA relative to total peptide was calculated and normalized to mean levels in the buffer control at t=~1/6 min.

c. A subpopulation of peptidyl-tRNA:80S complexes is refractory to Jlp2-mediated release over extended incubation periods. Extended time course (1 hour) Western blot analysis of *in vitro* reconstitution reactions using purified wild-type or H44A mutant Jlp2-FLAG and sucrose density gradient-enriched peptidyl-tRNA:80S complexes from *jlp2Δltn1Δrqc2Δ* cells.

d. 80S complexes used for *in vitro* reconstitution remain intact throughout the reaction. Analytical sucrose density gradient fractionation of *in vitro* reactions using high salt and physiological salt sucrose density gradient-enriched peptidyl-tRNA:80S from *jlp2Δltn1Δrqc2Δ* cells and purified Jlp2. $A_{260}$ profiles and Western blot analysis of resulting top (ribosome-free), 60S and 80S fractions are shown.

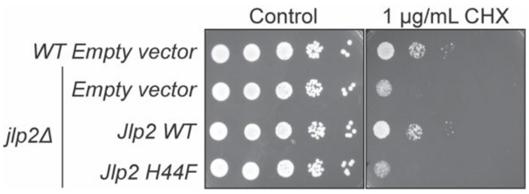


**Supplementary Figure S5. The release factor activity of Jlp2 is required for protection against translation elongation stress.**

Spot assay showing the growth of WT cells containing an empty vector and *jlp2Δ* cells containing either an empty vector, or overexpression vectors for WT or H44F Jlp2 spotted in serial dilutions on CHX-containing plates, imaged after 4 days and 8 days for control and 1 µg/mL respectively.
